# Soil depth governs microbial community assembly and enzymatic activity in extreme environments

**DOI:** 10.1101/2022.08.17.504275

**Authors:** Xin Jing, Aimée T. Classen, Daijiang Li, Litao Lin, Mingzhen Lu, Nathan J. Sanders, Yugang Wang, Wenting Feng

**Affiliations:** State Key Laboratory of Herbage Improvement and Grassland Agro-Ecosystems, and College of Pastoral Agriculture Science and Technology, Lanzhou University, Lanzhou, 730000, Gansu, China; Department of Ecology and Evolutionary Biology, University of Michigan, Ann Arbor, Michigan, USA; Department of Biological Sciences, Louisiana State University, Baton Rouge, Louisiana, USA; Center for Computation & Technology, Louisiana State University, Baton Rouge, Louisiana, USA; State Key Laboratory of Efficient Utilization of Arid and Semi-arid Arable Land in Northern China, the Institute of Agricultural Resource and Regional Planning, Chinese Academy of Agricultural Sciences, Beijing, 100081, China; Santa Fe Institute, Santa Fe, NM 87501, USA; State Key Laboratory of Desert and Oasis Ecology, Xinjiang Institute of Ecology and Geography, Chinese Academy of Sciences, Urumqi, Xinjiang, 830011, China; Fukang Station of Desert Ecology, Chinese Academy of Sciences, Fukang, Xinjiang, 831505, China; University of Chinese Academy of Sciences, Beijing, 100049, China; School of Grassland Science, Beijing Forestry University, Beijing, China

**Keywords:** ecosystem functioning, extreme environments, microbial beta-biodiversity, soil macroecology, structure-function relationships, Taklamakan desert

## Abstract

**Aim:** A fundamental challenge in soil macroecology is to understand how microbial community structure shapes ecosystem functions along environmental gradients of land surface (i.e., horizontal dimension). However, little is known of microbial community structure-function relationships along environmental gradients of soil depth (i.e., vertical dimension) in extreme environments. A full understanding of the consequences of environmental change for microbial communities structure and subsequent changes in microbial functions could enable more accurate predictions of extreme environmental change effects. Here, we leveraged a 200-km desert soil salinity gradient that is created by a 12-year saline-water irrigation to evaluate how soil microbial community structure-function relationships change with soil salinity in the horizontal and vertical dimensions.

**Location:** The Tarim basin of Taklamakan desert.

**Taxa:** Soil bacteria and fungi.

**Methods:** We assessed the prime ecological processes controlling the assembly of microbial communities and the activity of enzymes relevant to carbon, nitrogen, and phosphorus cycling along soil salinity gradients across study sites (horizontal dimension) and soil depths (vertical dimension) by using the general linear model, hierarchical variance partitioning, and path model.

**Results:** Differences in soil depth (on the scale of meters) was as important as geographic distance (on the scale of kilometers) in shaping the structure of bacterial and fungal communities, while both the vertical and horizontal variability in enzymatic activity were largely attributed to the increase in the heterogeneity of soil properties, such as soil texture, water content, and pH.

**Main conclusions:** Our results suggest that dispersal limitation and environmental heterogeneity, not soil salinization, along soil depth governs microbial community assembly and enzymatic activity, respectively. This work highlights that conservation efforts of soil macroecology should consider soil depth as a key attribute in the face of ongoing salinization in arid ecosystems.

## 1. Introduction

Understanding how soil microbial community structure, i.e., community biodiversity and composition, shapes spatial variability in ecosystem functions across environmental gradients is becoming a fundamental challenge in the field of soil macroecology (Chu, Gao, Ma, Fan, & Delgado-Baquerizo, 2020; Eisenhauer et al., 2021; White et al., 2020; X. F. Xu et al., 2020). Although soil microbial biodiversity is important for numerous ecosystem functions along environmental gradients on land surface (i.e., horizontal dimension) (Bardgett & van der Putten, 2014; Delgado-Baquerizo et al., 2020; X. Jing et al., 2015; van der Heijden, Bardgett, & van Straalen, 2008; D. H. Wall, Nielsen, & Six, 2015), it is a relatively new endeavor to investigate the influences of spatial variability in soil microbial community composition on spatial variability in microbial functions (Bardgett & van der Putten, 2014; T. Bell et al., 2009; Berlinches de Gea, Hautier, & Geisen, 2022; Xin Jing et al., 2022; Martinez-Almoyna et al., 2019; Mori et al., 2016; Peay, Kennedy, & Talbot, 2016; Talbot et al., 2014). One of the grand challenges is that hyper-diverse microorganisms inhabit soil matrix (Coleman & Whitman, 2005; Díaz & Malhi, 2022; Nielsen, Wall, & Six, 2015), as there can be 100-9000 microbial species/strains in a small soil sample (Bardgett & van der Putten, 2014) vs. 1∼32 species in a grassland biodiversity experiment (Hector et al., 1999). This makes it much difficult to uncover microbial community structure-function relationships (Peay et al., 2016).

One strategy to address this challenge is to study community structure-function in extreme environments, such as desert, glaciers, and hypersaline habitats, where environmental constraints are high and biological community structure is relatively simple (Ertekin, Meslier, Browning, Treadgold, & DiRuggiero, 2021; Osborne, Hall, Kronfeld-Schor, Thybert, & Haerty, 2020; Shu & Huang, 2022; Diana H Wall & Virginia, 1999). Moreover, the study of extreme environments provides not only useful information on how environmental conditions shape the geographic distributions of micro- and macro-organisms, but also unique opportunities to predict how community structure and functions responds to accelerated environmental change (N. Fierer et al., 2012; Shu & Huang, 2022; Diana H Wall & Virginia, 1999).

In extreme arid environments (e.g., desert), niche-based environmental filters such as salinity, soil water, and nutrient availability can shape the composition and function of microbial communities, but these drivers can also change with soil depth (i.e., vertical dimension) (Rath, Fierer, Murphy, & Rousk, 2019; Rath, Murphy, & Rousk, 2019). Soil depth is a spatial dimension that is often overlooked in the field of soil macroecology (Eisenhauer et al., 2021; White et al., 2020) but important for an explicit understanding of how microbial community structure relates to microbial functions through soil depth profiles (Noah Fierer, Schimel, & Holden, 2003). In fact, prior work highlights that soil depth influences the diversity of microbial communities, because important drivers, such as soil organic carbon, nutrient, salinity, oxygen concentration, and moisture, vary with soil depth (Noah Fierer et al., 2003; Powell et al., 2015; Stone, Kan, & Plante, 2015; Sun, Wang, Hui, Jing, & Feng, 2020). However, although soil depth also regulates microbial functions such as organic matter decomposition and nutrient cycling, we do not know much about how the vertical patterns of species turnover in microbial community composition, i.e., β-diversity, one of the important components of community structure, are related to the spatial variability in microbial functions along soil profiles.

Here, we address the question how microbial community structure affects the spatial variation in microbial activity following a 12-year saline-water irrigation (i.e., soil salinization) in the Taklamakan desert, the world’s 2^nd^ largest shifting sand desert with the area of roughly the size of Germany (Fig. S1). The potential consequences of soil salinization on microbial physiology, growth and community structure have been extensively studied (Rath & Rousk, 2015), but long-term impacts of salinization on soil microbial community structure and enzymatic activity in both the horizontal and vertical dimensions have been rarely investigated. We, therefore, proposed a spatial dimension partitioning approach (Fig. 1) to test two hypotheses: soil salinization will enhance soil microbial species turnover (Hypothesis 1) and mediate the microbial community structure-enzymatic activity relationships (Hypothesis 2) in the horizontal and vertical spatial dimensions. For hypothesis 2, we specifically predict that soil salinization will enhance species turnover when controlling for other environmental factors, e.g., geographic distance, differences in soil chemical properties and nutrient pools (H2a); soil salinization will indirectly enhance enzymatic turnover via soil bacterial and fungal species turnover (H2b). Using the analytical framework, we evaluate soil bacterial and fungal community structure and enzymatic activity relevant to soil carbon, nitrogen, and phosphorus cycling along a unique 200-km salinity gradient (i.e., the horizontal dimension; Fig. S2a and S2b). At each site we also examine bacterial and fungal community assembly and enzymatic activity across a one-meter soil profile (i.e., the vertical dimension; Fig. S2a and S2c), an overlooked dimension of soil macroecology. In addition to soil salinity, we assess a range of ecological factors, i.e., soil physical and chemical properties and nutrient pools (Table S1) that could mediate microbial community structure-enzymatic activity relationships.

**Fig. 1.**
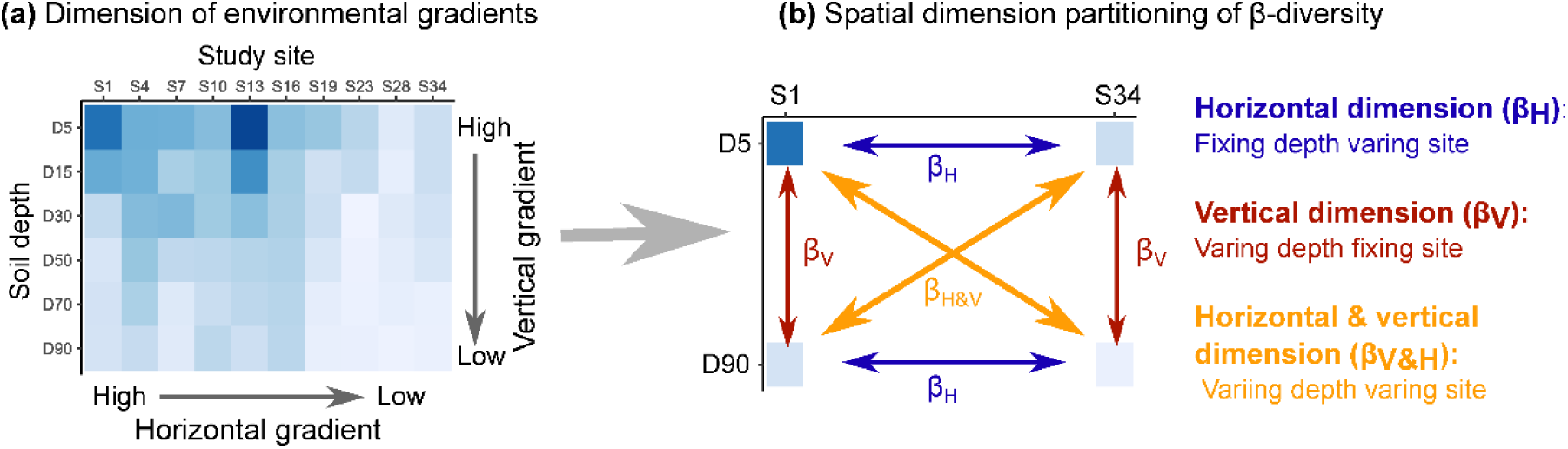
Conceptual depictions of spatial dimension partitioning approach. (a) heatmap denotes the horizontal and vertical dimension of environmental gradients, i.e., soil salinity. The gradients of soil salinity were created by a 12-year saline-water irrigation in the Taklamakan desert. (b) a framework for illustrating the spatial dimension partitioning of β-diversity. β_H_, β_V_ and β_H&V_ denotes species turnover along the horizontal, vertical, and horizontal & vertical gradient of soil salinity, respectively.

## 2. Materials and Methods

### 2.1. Study sites and data collection

We conducted this study in shrubland vegetation located along the Tarim desert highway, which crosses the Tarim basin of Taklamakan desert (37.1−41.8° N, 82.6−84.3° E) (Fig. S1). The Taklamakan desert is arid, with a mean annual precipitation of 24.6 mm yr^-1^ and a mean annual potential evaporation of 3639 mm yr^-1^. The mean annual air temperature is 2.4°C, with the coldest monthly mean temperature reaching −8.1°C in December and the warmest monthly mean temperature reaching 28.2°C in July. Most of the desert is bare land without vegetation cover, except for a few extremely drought- and salt-tolerant shrub species, such as, *Tamarix taklamakanensis* and *Calligonum taklamakanensis*. Soils in the desert are aeolian sandy soil. The top 0 20 cm soil is composed of 87.3% of sand, 12.4% of silt, and 0.3% of clay (J. Zhang et al., 2016). The soils have an extremely low organic matter content (0.74 g kg^-1^) and total nitrogen (0.14 g kg^-1^) in the top 0 10 cm soil depth (Li, Lei, Zhao, Xu, & Li, 2015).

In 2005, the shrubland vegetation was established to prevent shifting sand dunes from damaging the Tarim desert highway. After 12 years of vegetation development, the plant community has stabilized, with *Haloxylon ammodendron* being dominant the community and *Calligonum aborescens*, *Tamarix ramosissima*, *Nitraria tangutorum*, *Populus euphratica* and *Elaeagnus angustifolia* with much lower abundance. The aboveground plant community did not vary along the 200 km salinity gradient of study region. Additionally, the aboveground biomass of dominant species *H. ammodendron* did not vary across the studied sites, probably leading to comparable amount of carbon inputs into the soil. The groundwater level in the Taklamakan desert ranges from 3 to 5 m, and the groundwater is charged by water from the Tianshan and Kunlun mountains (Fig. S1). The groundwater was pumped for drip irrigation to water shrubs in the vegetation. The drip irrigation pipes run along the rows of shrubs, and the vegetation was irrigated twice per month from March to May and from September to October, and three times per month from June to August with ∼30 L of water per plant per irrigation event (X. Xu, Li, & Wang, 2006).

We surveyed ten shrubland sites along an extreme gradient of irrigation water salinity using a regression design of field experiment (see Pelini et al., 2011 for an example of the regression design). The regression design used a gradient of water salinity that varied from 3.2 to 38.0 g L^-1^ of total dissolved salts (the normal range of seawater salinity is between 33-37 g L^-1^ of dissolved salts) from the south to the north sampling site (Figs. S1 and S2b). We collected irrigation water and soil samples in early October 2017, which is the end of the growing season in the desert. At each study site, three replicates of irrigation water were collected from running water that pumped from wells. Irrigation water samples were stored in 500 ml plastic bottles, sealed, and taken to the laboratory for water salinity analysis. We sampled soils using soil cores (5 cm in diameter) at three randomly selected plots, which were located between the two rows of shrubs and 30 50 cm away from a shrub. We collected soils to the depth of 100 cm with sampling intervals of 20 cm, except that we segmented the topsoils to 0 10 and 10 20 cm. We collected 180 soil samples (10 sites × 6 soil layers × 3 replicates). Using plastic bags and coolers, a subsample (∼30 g) of each soil was kept cool and immediately returned to the laboratory. We sieved all soils to 2 mm and removed rocks and roots by hand. We stored soils at 20°C for microbial sequencing and at 4°C for the assays of soil enzymatic activity. We air-dried the remaining soils to analyze soil properties, including salinity, pH, water content, texture, total organic carbon, total nitrogen, and available phosphorus.

We analyzed the salinity of irrigation water in the following ways. We determined the concentrations of Na^+^, K^+^, Ca^2+^, and Mg^2+^ using a spectrometer (iCAP 6300-ICP-OES) with a CID detector (Thermo Scientific, MA, USA). Using titration, we analyzed the concentration of Cl^-^ with AgNO_3_, SO_4_ ^2-^ with ethylenediaminetetraacetic acid (EDTA), and CO_3_ ^2-^ and HCO_3_ ^-^ with sulfuric acid until all bicarbonate was neutralized. Next, we calculated the salinity of irrigation water as the sum of the concentrations of Na^+^, K^+^, Ca^2+^, Mg^2+^, Cl^-^, SO_4_ ^2-^, CO_3_ ^2-^ and HCO_3_ ^-^.

We measured soil salinity and pH in a 1:5 suspension of soil to water. We analyzed soil salinity using the methods described above for irrigation water salinity. We analyzed soil pH using the Mettler Toledo FiveEasy FE28 (Greifensee, Switzerland). We measured soil gravimetric water content by oven drying 10 g of fresh soils at 105°C for 24 h. We measured soil texture using the laser diffraction method (Sympatec GmbH, System-PartikelTechnik, Clausthal-Zellerfeld, Germany) after removing the organic material by 10% hydrogen peroxide and removing carbonates by 0.2% hydrochloric acid. We analyzed soil organic carbon using the Walkley and Black method of oxidizing organic carbon with K_2_Cr_2_O_7_-H_2_SO_4_ (Nelson & Sommers, 1983). We analyzed soil total nitrogen using an AutoKjeldahl Unit model K370 (BUCHI, Flawil, Switzerland) and soil available phosphorus using the sodium bicarbonate extraction with Mo-Sb Anti-spectrophotometer method.

To analyze soil bacterial and fungal diversity, we extracted soil DNA using the PowerSoil DNA extraction kit (Qiagen, Carlsbad, CA, USA) following the manufacturer’s protocol. We amplified the V4 region of 16S rRNA gene for soil bacteria using the primer pairs 515F/806R and the ITS1 region for soil fungi using the primer pairs ITS5-1737F/ITS2-2043R. We performed PCR using the BioRad S1000 thermal cycler (Bio-Rad Laboratory, CA, USA) after mixing 25 µl of 2x Premix Taq polymerase, 2 µl of 10 mM primer pairs, 3 µl of 20 ng µl^-1^ DNA template and 20 µl nuclease-free water. Next, we ran the thermal cycling as follows: initial denaturation at 94°C for 5 min, followed by 30 cycles of denaturation at 94°C for 30 s, annealing at 52°C for 30 s, and elongation at 72°C for 30 s, with a final step of 72°C for 10 min. We repeated this PCR procedure three times for each sample. We next mixed the PCR products using GeneTools Analysis Software (Version 4.03.05.1, SynGene). The process was followed by purifying the DNA products using the EZNA Gel extraction kit (Omega Bio-Tek, GA, USA). Following the manufacturer’s instruction, we generated sequencing libraries using the NEBNext ® Ultra™ DNA Library Prep Kit for Illumina® (New England Biolabs, MA, USA) and sequenced the libraries on an Illumina HiSeq2500 platform (Illumina PE250, CA, USA).

We processed raw DNA sequences using the Trimmomatic software (V0.33, http://www.usadellab.org/cms/?page=trimmomatic). Specifically, we excluded the raw DNA sequences by removing paired-end reads with N, read quality <20, and read length <100 bp. Next, we allocated the reads on the paired end to the corresponding soil samples using the Mothur pipeline (V1.35.1, http://www.mothur.org). We removed the barcode and primers to derive the paired-end clean reads, which were merged with FLASH (V1.2.11, https://ccb.jhu.edu/software/FLASH). The minimum length of overlap was 10 bp and the maximum ratio of number of mismatches to overlap length was 0.1. Next, we assigned these clean tags to OTUs with 97% similarity using USEARCH software (V8.0.1517, http://www.drive5.com/usearch). We used representative sequences for taxonomic classification using the RDP classifier according to the Greengenes database (http://greengenes.secondgenome.com) for bacteria and the Unite database (http://unite.ut.ee/index.php) for fungi in the QIIME pipeline (version 2; http://qiime.org/scripts/assign_taxonomy.html).

To explore microbial functions, we assayed the potential activities of four hydrolytic enzymes (β-1, 4-glucosidase, BG; β-_D_-cellobiohydrolase, CB; β-1, 4-N-acetyl-glucosaminidase, NAG; alkaline phosphatase, ALP) using the 96-well microplate and the fluorometric technique (C. W. Bell et al., 2013; German et al., 2011). We selected these enzymes because their potential activities are commonly measured in soils and used as proxies for the cycling of carbon (BG and CB), nitrogen (NAG), and phosphorus (ALP) (Sinsabaugh et al., 2008). Briefly, sieved fresh soil (1.5 g) was suspended in 150 ml of 50 mM Tris buffer (pH = 8.0). Soil slurries (200 μl) of each sample were transferred to eight wells of the 96-well microplate and mixed with 50 μl of 200 μM standard fluorometric substrates of each enzyme. The microplates were incubated for 2.5 h in the dark at 25°C. The quantity of fluorescence was determined at 360 nm excitation and 460 nm emission in a 96-well microplate reader (Biotek Synergy 2, Winooski, VT, USA). A multiple-point calibration curve was used to calculate enzymatic activities (C. W. Bell et al., 2013). The units were expressed as nmol g dry weight^−1^ h^−1^.

### 2.2. Analyses and Statistics

We used a two-step approach to examine how soil salinization affects soil microbial community structure and enzymatic activity.

Step 1, to test whether increasing soil salinity enhances species turnover (Hypothesis 1), we developed a dimension partitioning approach (Khattar, Macedo, Monteiro, & Peres Neto, 2021) to quantify the impacts of vertical and horizontal differences in soil salinity on spatial variability in the structure of bacterial and fungal communities. We calculated the pair-wise dissimilarity matrix between soil microbial assemblages for all combinations of study sites and soil layers (*n* = 60). The lower triangle of the pair-wise dissimilarity matrix was converted into a single vector, resulting in 1770 observations (60 × 59 / 2). We further partitioned the single vector into three spatial dimensions, including the horizontal, vertical, and horizontal & vertical dimensions. Differences in community structure of the same soil layer between study sites were defined as the horizontal dimension (10 × 9 / 2 × 6 = 270 observations), between soil layers within a study site were defined as the vertical dimension (6 × 5 / 2 × 10 = 150 observations), and between soil layers and between sampling sites were defined as the horizontal & vertical dimension (1350 observations).

To measure the spatial variability in soil microbial community structure between assemblages, we used the Podani family of Sorensen indices (Legendre, 2014; Podani & Schmera, 2011; Schmera, Podani, & Legendre, 2020), including the Sorensen dissimilarity index (*β_sor_*), species replacement (*β_repl_*), and richness difference (*β_rich_*). The Sorensen dissimilarity index quantifies the total differences in species composition between assemblages. It was partitioned into two additive components, i.e., species replacement and richness difference. The former quantifies the species in one assemblage that are replaced by different species in another assemblage, and the latter estimates differences in the number of species between assemblages. We used the following equations:

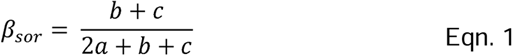

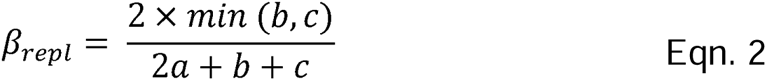

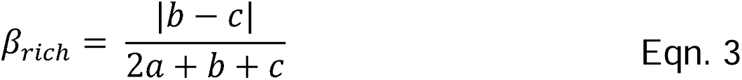

where *a* denotes the number of species present in both assemblages, *b* denotes the number of species present in assemblage 1 not in assemblage 2, and *c* denotes the number of species present in assemblage 2 not in assemblage 1.

To compare the species turnover rate with differences in soil salinity, we calculated the Euclidean distances as the measure of differences in soil salinity. We calculated differences in soil salinity using the concentrations of soil Na^+^, K^+^, Ca^2+^, Mg^2+^, Cl^-^, SO_4_ ^2-^, and HCO_3_ ^2-^ as follows:

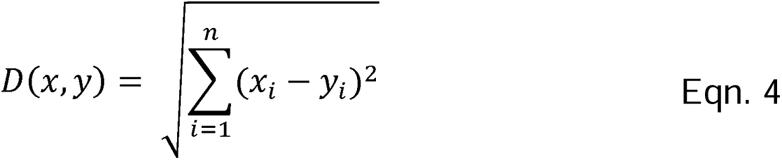

where *D(x, y)* denotes the Euclidean distance, *n* is the number of salt ions (*n* = 7), *x_i_* and *y_i_* denote the concentration of salt ions in the sample *x* and sample *y*, respectively. We regressed pair-wise compositional dissimilarity (i.e., β-diversity) against differences in soil salinity across the horizontal and vertical spatial dimensions. We used the standardized regression slopes to compare the rate of species turnover among spatial dimensions and the permutation test to examine the significance of regression slopes.

Step 2, to test whether increasing soil salinity mediate microbial community structure-enzymatic activity relationships (Hypothesis 2), we used hierarchical variance partitioning and path model. We conducted hierarchical variance partitioning to estimate the proportion of spatial variability in the structure of microbial communities (Hypothesis 2a). We used the Euclidean distances to calculate geographic distance and differences in soil depth, salinity, properties, and nutrient pools (Table S1). Specifically, the geographic distance was calculated using the geographic coordinates of the sampling sites. Differences in soil depth were calculated using the mean depth of each sampling layer. Differences in soil property were calculated using soil clay content, silt content, sand content, pH, and moisture. Differences in nutrient pools were calculated using the data of soil organic carbon, soil total nitrogen, and available phosphorus (Xin Jing et al., 2022). All combinations of explanatory variables were fitted into regression models following the hierarchical partitioning algorithm of Chevan and Sutherland (1991). The independent effects of each explanatory variable were determined based on the goodness-of-fit measures (i.e., R^2^) of regression models, with the target variables (microbial community structure or enzymatic activity) as the dependent variable. We used 9,999 bootstrapped sampling iterations to calculate the 95% confidence interval of the independent effects of each explanatory variable.

We used the path model to estimate the indirect effects of soil salinity on enzymatic turnover via bacterial and fungal species turnover (Hypothesis 2b). We considered the potential enzymatic activities of BG, CB, NAG and ALP as a proxy for microbial functions (Burns et al., 2013; Fanin et al., 2022; Talbot et al., 2014). We used the Euclidean distances as the measures of spatial variability in microbial functions, i.e., enzymatic turnover or differences in enzymatic activity (Xin Jing et al., 2021; Xin Jing et al., 2022; Martinez-Almoyna et al., 2019; Mori et al., 2016). The path model was also used to assess the relative importance of dispersal limitation and environmental filtering on the structure of soil microbial communities (Xin Jing et al., 2021; Xin Jing et al., 2022). For soil bacterial and fungal community structure, we considered geographic distance and differences in soil depth as the proxies for dispersal limitation (Powell et al., 2015) and differences in soil salinity, properties, and nutrient pools as the proxies for environmental filtering (Xin Jing et al., 2022; Martiny et al., 2006). Using path model, we also estimated the relative importance of environmental variables and community structure of soil bacteria and fungi on differences in microbial enzymatic activity. We used the P value of χ^2^ statistic test (P >0.05), comparative fit index (CFI >0.90), root mean square error of approximation (RMSEA <0.05), and standardized root mean square residual (SRMR <0.10) as measures of path model evaluation and selection (Grace, 2020). Since a large sample size (*n* > 100) can lead to the P value of χ^2^statistic test that was less than 0.05 (Grace, 2020), we only used CFI, RMSEA, and SRMR to evaluate and select models for overall and horizontal and vertical spatial dimensions. We used 9,999 bootstrapped sampling iterations to test the significance of the path coefficients. All path models fitted the data well (Table S2).

The dimension partitioning was carried out for all the variables, including soil microbial community structure and enzymatic activity, geographic distance, depth difference, salinity difference, property difference, and nutrient pool difference. Prior to data analyses, we removed singleton OTUs, which are present only once across all microbial assemblages, from soil microbial community data and pooled all the data to the level of site × soil layer to remove the non-independent influences of sample replicates. All statistical analyses were performed in R version 3.6.1 (R Development Core Team, 2019) using the packages ‘lmPerm’ (Wheeler, Torchiano, & Torchiano, 2016), ‘hier.part’ (Nally & Walsh, 2004), and ‘lavaan’ (Rosseel, 2012).

## 3. Results

Bacterial total β-diversity (β_sor_) increased with differences in soil salinity across horizontal, vertical, and horizontal & vertical spatial dimensions (Fig. 2a, 2c, and 2e). Bacterial richness difference (β_rich_) showed weakly positive relations with differences in soil salinity, while bacterial replacement was low at high level of differences in soil salinity across the horizontal and horizontal & vertical spatial dimensions (Fig. 2a and 2e). Fungal β_sor_ decreased with differences in soil salinity in the horizontal spatial dimension (Fig. 2b). Fungal β_repl_ decreased with differences in soil salinity in the horizontal spatial dimension but increased with differences in soil salinity in the vertical spatial dimension (Fig. 2b and 2d). Fungal β_rich_ increased with differences in soil salinity in the horizontal spatial dimension, while had no relations with differences in soil salinity in other spatial dimensions (Fig. 2b, 2d and 2f).

**Fig. 2.**
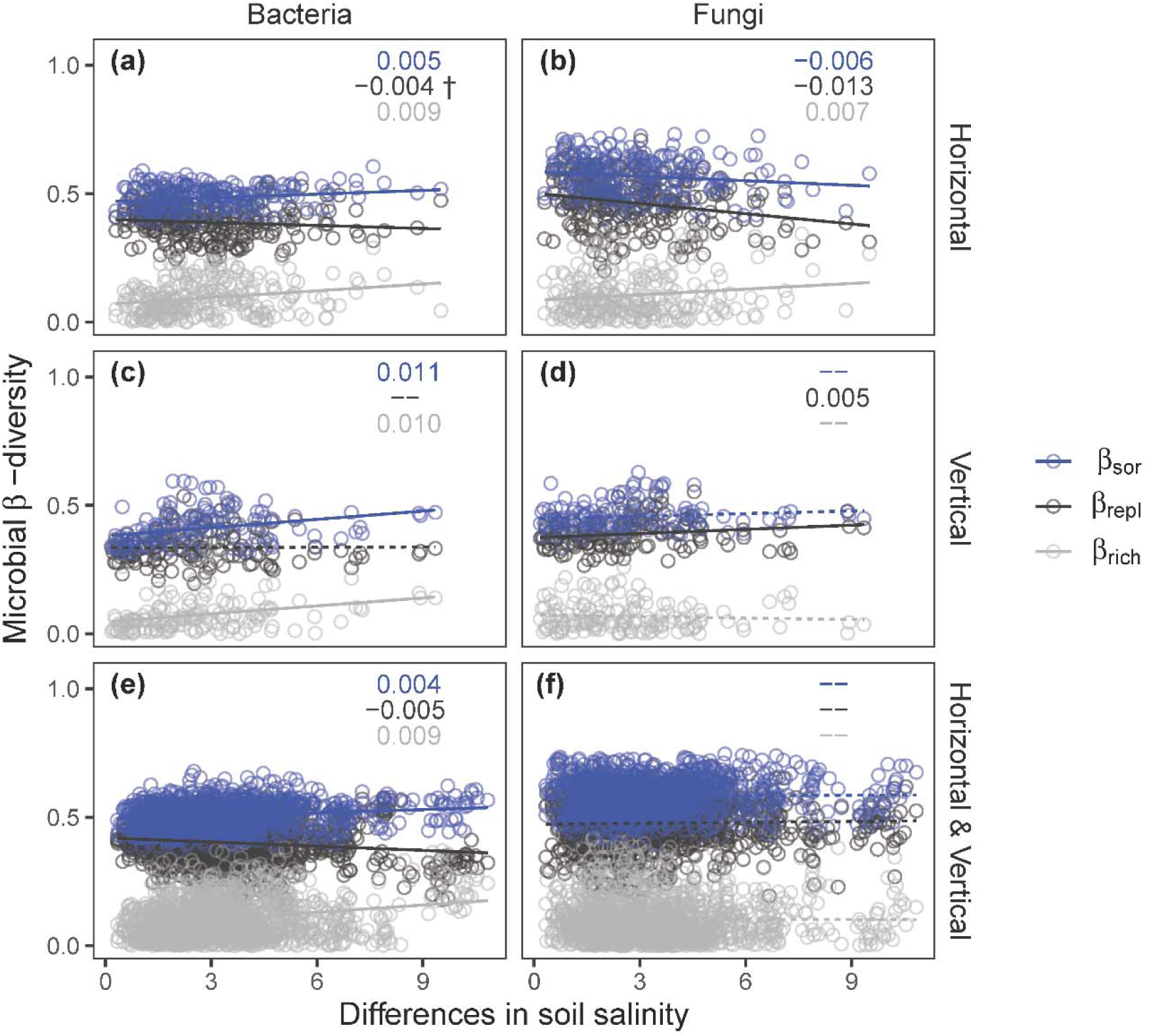
Relationships between differences in microbial β-diversity and soil salinity across the horizontal and vertical spatial dimensions. Solid lines denote significant regressions (if †, 0.05 < P < 0.10, otherwise P < 0.05) and dashed lines denote nonsignificant regressions (P > 0.10). Significant standardized regression coefficients are shown at the top right side of each panel and nonsignificant standardized regression coefficients are shown as ‘--’. β_sor_ = Sorensen dissimilarity index (blue); β_repl_= replacement component of β_sor_(dark gray); β_rich_ = richness difference component of β_sor_(gray).

Differences in soil salinity did not exert significant independent effects on soil microbial β_sor_, β_repl_, and β_rich_ (Figs. 3, S3 and S4; Table S3). By contrast, geographic distance and differences in soil depth had the largest independent effects on differences in soil bacterial and fungal β_sor_ in the horizontal or vertical spatial dimensions (Fig. 3a-3f). Geographic distance explained 20.7-25.2% of the variance in differences in soil bacterial β_sor_and 24.1-25.3% of the variance in differences in soil fungal β_sor_ (Fig. 3a, 3c, 3d and 3f; Table S3). Throughout the vertical dimension, the differences in soil depth explained 21.9% (95% confidence interval [12.4, 32.7]) of the variance in differences in soil bacterial β_sor_ (Fig. 3b Table S4) and 23.7% [12.7, 35.8] of the variance in differences in soil fungal community structure (Fig. 3b and 3e; Table S3).

**Fig. 3.**
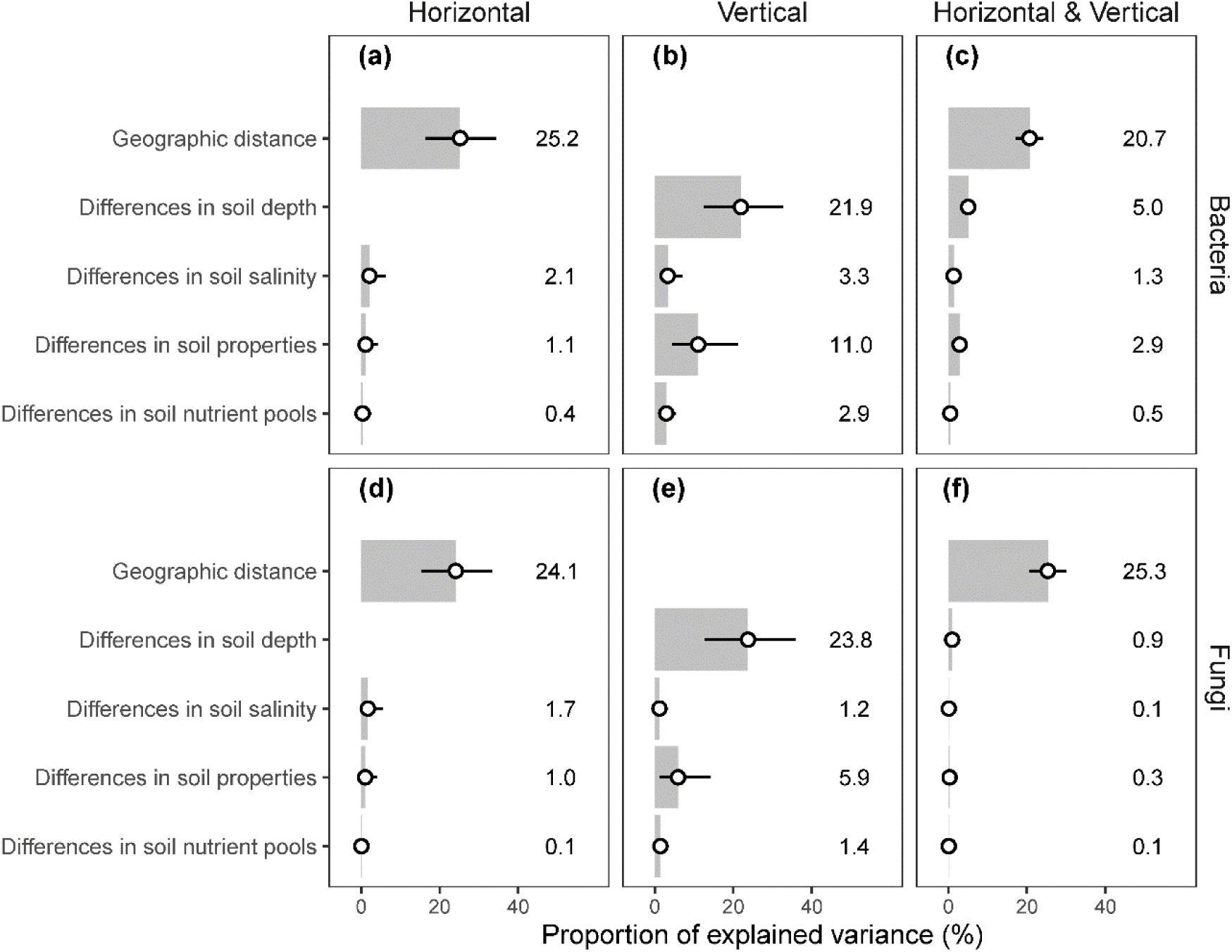
Independent effects of explanatory variables on microbial β-diversity across the horizontal and vertical spatial dimensions by hierarchical variance partitioning. Bars represent the pure variance explained by the explanatory variables, and the proportion of explained variance is shown along the bars. The 95% confidence interval is given. We show only the Sorensen dissimilarity index, see Figs. S3-S4 and Table S2 for the summary of the replacement and richness difference component of Sorensen dissimilarity index.

Path analysis showed that geographic distance and differences in soil depth, salinity, properties, and nutrient pools separately explained 14% 27% of the variance in differences in enzymatic activity, 29% 39% of the variance in differences in soil bacterial community structure, and 27% 32% of the variance in differences in soil fungal community structure (Fig. 4a-4c). However, differences in enzymatic activity were not explained by differences in soil microbial community structure (standardized path coefficients *β_std_* < 0.13 in all spatial dimensions; Figs. 4 and S5-S10; Table S4). Differences in soil properties but not soil salinity had the largest effects on differences in enzymatic activity in the vertical dimension, which were as important as the effects in the horizontal dimension (Fig. 4). In addition, differences in soil bacterial and fungal community structure were weakly associated with differences in soil salinity (Fig. 4). The effects of differences in soil depth (Fig. 4b) on soil bacterial and fungal β_sor_ along the vertical dimension were as important as the effects of geographic distance (Fig. 4a and 4c) along the horizontal dimension on soil bacterial and fungal β_sor_.

**Fig. 4.**
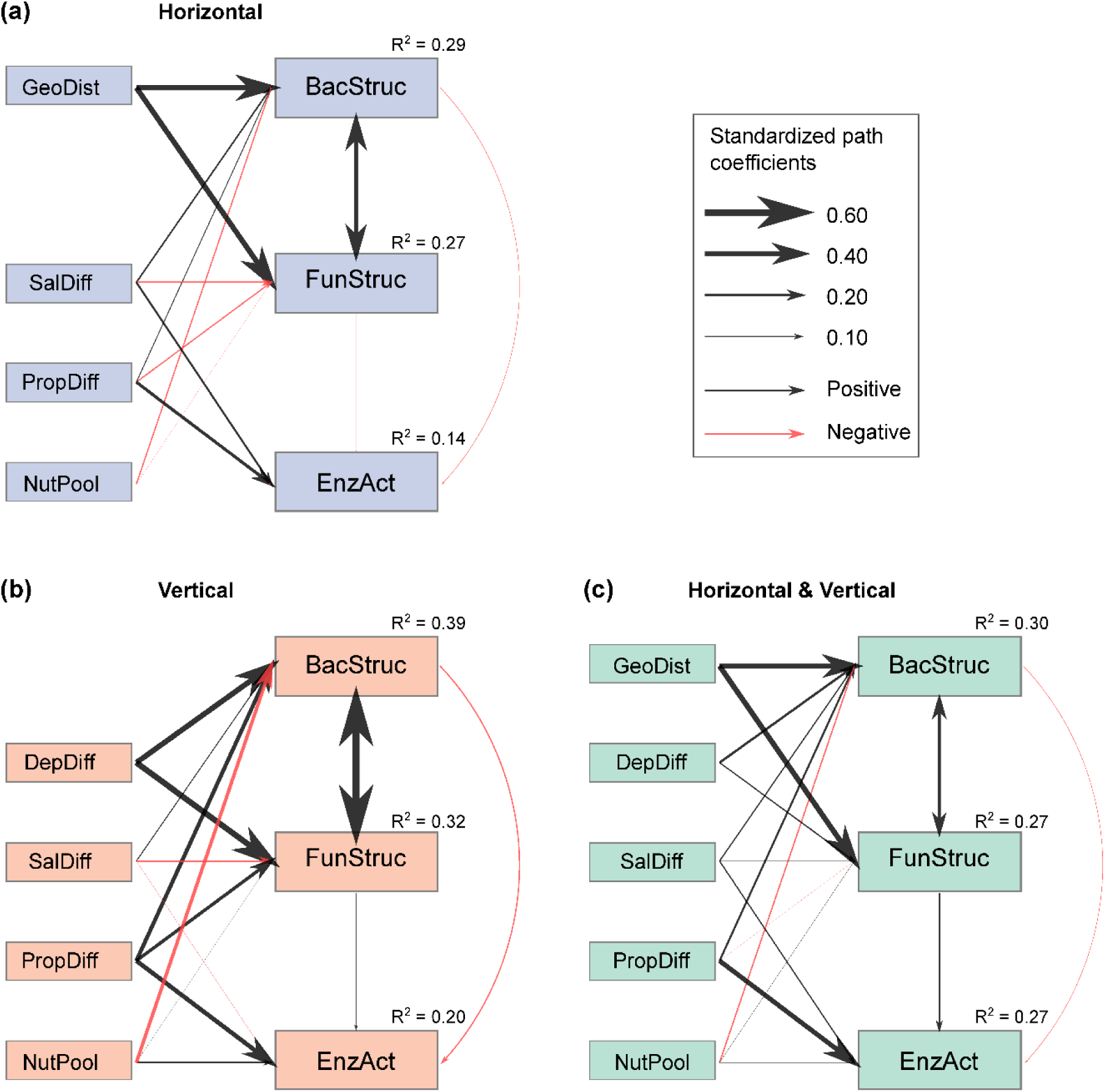
Path model shows the direct and indirect effects of soil salinity on microbial community structure and enzymatic activity across the horizontal and vertical spatial dimensions. Three path models are shown based on spatial dimensions, including a) pure horizontal dimension; b) pure vertical dimension; c) combined horizontal & vertical dimension. The width of the arrows is proportional to the standardized path coefficients. Black arrows denote positive path coefficients and red arrows denote negative path coefficients. Bootstraps are used for the significance test of path coefficients (see Tables S3-S4 for details). R^2^ denotes the variance in community structure and enzymatic activity explained by environmental factors, including geographic distance (GeoDist), differences in soil depth (DepDiff), differences in soil salinity (SalDiff), differences in soil properties (PropDiff), and differences in nutrient pools (NutPool). BacStruc, FunStruc, and EnzAct denote differences in bacterial community structure, fungal community structure, and enzymatic activity, respectively. We show only the Sorensen dissimilarity index, see Figs. S10-S11 and Tables S3-S4 for summary of the replacement and richness difference component of Sorensen dissimilarity index.

## 4. Discussion

Here, we used a unique environmental (i.e., salinity) gradient that display spatial variation in both the horizonal and vertical dimensions. Using this salinity gradient, we were able to separate the influences of horizontal soil salinization (on the scale of kilometers) on microbial community assembly and enzymatic activity from the influences of vertical soil salinization (one the scale of meters), which is an overlooked dimension of soil macroecology. Our first hypothesis that increasing soil salinity will lead to enhanced microbial species turnover was weakly supported by fungal species turnover in the vertical spatial dimension. Instead, we found that richness difference contributed to variation in bacterial and fungal species composition in both the horizontal and vertical spatial dimensions (Fig. 2). Our second hypothesis was rejected because geographic distance in the horizontal dimension and difference in soil depth in the vertical dimension, rather than differences in soil salinity, consistently determined variation in bacterial and fungal species composition (Fig. 3). Differences in soil properties rather than soil salinity enhanced the spatial turnover in enzymatic activity (Fig. 4). Our findings suggest that soil depth serves as a test bed for predicting changes in soil microbial community structure and enzymatic activity in extreme environments.

### 4.1. Soil depth as a new frontier for studies in soil macroecology

In this work, we found that spatial variability in the structure of soil microbial communities was strongly associated with geographic distance and differences in soil depth rather than differences in soil salinity. This result suggests that soil bacterial and fungal communities are resistant to changes in soil salinity but show a dissimilar community structure with an increase in spatial distance both horizontally and vertically along soil depth profile. Our finding that soil microbial community structure is resistant to changes in soil salinity is in contrast with previous work (Rath, Fierer, et al., 2019; Van Horn et al., 2014; K. Zhang et al., 2019). One reason must be that soil salinity is extremely high after 12-year of saline-water irrigation, which created a long-lasting harsh environmental condition (Schradin, Makuya, Pillay, & Rimbach, 2023). Soil microbes may have adapted well to the harsh environment, as low species turnover was associated with differences in soil salinity (Fig. 2). However, our work indicates that soil depth is as important as geographic distance in shaping the structure of bacterial and fungal community in the saline desert soil (Figs. 3 and 4). The importance of soil depth is supported by an early study showing that random assembly processes of soil microbial communities (e.g., dispersal limitation) intensified with soil depth because soil microorganisms disperse passively (Powell et al., 2015). Our results indicate that soil depth and geographic distance are equally important when the overall spatial variability in microbial structure is partitioned into the pure vertical and horizontal spatial dimensions (Fig. 3a,b). Without spatial dimension partitioning, we will underestimate the contribution of soil depth to the assembly processes of soil microbial communities. Together, our work highlights that dispersal limitation in soil microbes is particularly important to the assembly of microbial communities in the vertical and horizontal dimensions of soil depth profiles in extreme environments. To maintain the essential and irreplaceable ecosystem functions and services provided by belowground biodiversity in extreme conditions or severely disturbed conditions, a workable plan is to preserve habitats along both the horizontal and vertical habitats along soil depth profiles.

Our study further indicates that the spatial variability in soil enzymatic activity is best explained by the heterogeneity in soil texture, pH, and water rather than differences in soil salinity and nutrient pools across soil depths. In extreme environments, environmental filters (e.g., salt stress) and resource levels (e.g., soil organic matter and microbial biomass) are expected to be the drivers of spatial variability in soil microbial community structure and function (Martiny et al., 2006; Rath, Fierer, et al., 2019; Rath, Murphy, et al., 2019; Rath & Rousk, 2015). On the contrary, our results suggest that differences in soil salinity and nutrient pools do not explain the activity of carbon, nitrogen, and phosphorus cycling related enzymes. It could be that soil salinity is too high in our study system and the priority of soil microbes is to survive rather than to thrive in such extreme saline environments, which does not allow soil microbes to produce energy-expensive enzymes (Rath & Rousk, 2015). As a result, soil salinity and resources are not limiting factors of enzymatic activity, and changes in soil salinity and nutrient pools did not alter enzymatic activity. However, our result indicates that reduced heterogeneity in soil properties, which could be induced by land degradation, soil erosion, and desertification, might result in loss of soil microbial activity and soil functionality, soil fertility, and plant growth. Since the variability in soil physical and chemical properties is created by human activities and nature ubiquitously, we should analyze the overlooked dimension of soil macroecology in the vertical dimension along soil depth profiles, which represent a valuable step towards soil conservation efforts in dealing with ongoing desertification in arid ecosystems.

### 4.2. Uncoupled microbial community structure-function relationships

The most salient result of this study is that differences in soil microbial community structure did not lead to differences in enzymatic activity in the hyper-saline environments. This result runs counter to previous work exploring these relationships in microbial systems with a focus on surface soils (Delgado-Baquerizo et al., 2020; Xin Jing et al., 2022; Mori et al., 2016). The uncoupled relationships between microbial structure and function were further verified across the horizontal and vertical dimensions of soil depth profile. Thus, our findings support functional redundancy (Coleman & Whitman, 2005; Peay et al., 2016), implying that even in extreme environments the structure of soil microbial communities differ, but their function in the same manner. As a result, the losses/gains of certain species may have limited effects on the spatial variability in enzymatic activity. This means that functionally redundant communities may safeguard ecosystem functions against the negative impacts of extreme environmental changes (Allison & Martiny, 2008).

There are two explanations for the uncoupled structure-function relationships under soil salinization. First, soil bacterial and fungal communities have likely adapted to the extreme conditions in desert, where the abiotic stress is high (Jansson & Hofmockel, 2020). As the soil becomes saltier and drier, soil microorganisms become inactive because the availability of carbon and nutrients in the soil decreases. In addition, soil microorganisms tend to minimize their activity, even becoming dormant, to conserve energy under harsh conditions. In either case, we would observe the uncoupled relationships between soil microbial community structure and enzymatic activity. Second, the uncoupled structure-function relationship is likely due to the mismatch of dominant driving factors responsible for microbial community structure and enzymatic activity. Our results show that geographic distance and differences in soil depth have the strongest independent effects on the structure of microbial communities, while differences in soil properties have the strongest independent effects on enzymatic activity. Different driving factors could have disproportionate influences on microbial community structure and enzymatic activity in soil, leading to uncoupled structure-function relationships.

In recent years, soil macroecologists have focused on microbial communities in surface soils because they have high biomass density and nutrient cycling activities (X. F. Xu et al., 2020). This study is unique in that it provides a new approach to assess microbial community structure-function relationships both horizontally and vertically along soil depth profiles with the same range of salinity variability in a typical extreme biome, i.e., the Taklamakan desert. However, large-scale studies remain the key to verify whether the findings hold true in other biomes, which may have a wider range of horizontal and vertical gradients that are induced by human activities or by nature under the context of soil degradation, e.g., soil salinization, acidification, and erosion. Further, it is important to note that the measure of four hydrolytic enzyme activities capture only a few microbial functions that are relevant to carbon, nitrogen, and phosphorus cycling. Future work that uses shotgun metagenomics could quantify broad shifts in microbial community functionality, e.g., the functions related to organotrophs and autotrophs (both by phototrophs and chemolithotrophs), advancing our understanding of the structure-function relationships of microbial communities in xeric or arid soils.

## 5. Conclusions

We find that environmental gradients along soil depth profile is as important as horizontal geographic distance in shaping microbial community structure and enzymatic activity. These findings highlight the need to analyze the overlooked dimension of soil macroecology, i.e., the vertical spatial dimension (Eisenhauer et al., 2021; White et al., 2020). Such information is crucial to predict the influences of global change drivers on real-world community structure-function relationships across horizontal and vertical spatial scales. Moreover, this study highlights that given the increase in desertification in arid ecosystems, soil conservation efforts should take soil depth into account as a critical factor governing the assembly of microbial communities and the activity of enzymes.

## Acknowledgments

This work was supported by the National Key Research and Development Program (2021YFE0114500), the National Natural Science Foundation of China (42077023, U1803342, 41930761), the K. C. Wong Education Foundation (GJTD-2020-14), and the Agricultural Science and Technology Innovation Program (ASTIP).

## Author Contributions

XJ, WF and YW designed the study. YW and WF conducted the field- and lab-work. XJ analyzed the data. XJ and WF wrote the first draft of the manuscript. All authors contributed to the conceptualization, review and editing of the manuscript.

## Data Availability

Sequence data were deposited in the European Nucleotide Archive with accession number PRJEB52395 (http://www.ebi.ac.uk/ena/data/view/OW744067-OW763241) for soil bacteria, and with accession number PRJEB46878 (http://www.ebi.ac.uk/ena/data/view/OU495976-OU501470) for soil fungi. The data and code will be publicly available in the Zenodo repository prior to publication.

## Biosketch

Xin Jing is a professor in the College of Pastoral Agriculture Science and Technology at Lanzhou University. He is interested in understanding the causes of biodiversity change and its consequences for ecosystem structure and function.

