## Supplementary for "Soil depth governs microbial community assembly and enzymatic activity in extreme environments"

**This file includes:**

Figures S1 to S10

Tables S1 to S4

**Figure captions**

**Fig. S1.** Geographic location of the study sites

**Fig. S2.** Horizontal and vertical salinity gradients generated by saline water irrigation

**Fig. S3.** Independent effects of explanatory variables on the replacement component of microbial β-diversity across the horizontal and vertical spatial dimensions

**Fig. S4.** Independent effects of explanatory variables on the richness difference component of microbial β-diversity across the horizontal and vertical spatial dimensions

**Fig. S5.** Bivariate relationships between differences in soil microbial community structure and differences in enzymatic activity across spatial dimensions

**Fig. S6.** Bivariate relationships between differences in soil microbial community structure and differences in enzymatic activity

**Fig. S7.** Bivariate relationships between differences in the replacement component of soil microbial community structure and differences in enzymatic activity

**Fig. S8.** Bivariate relationships between differences in the richness difference component of soil microbial community structure and differences in enzymatic activity

**Fig. S9.** Path models for testing whether increasing soil salinity indirectly enzymatic turnover via the replacement component of Sorensen dissimilarity

**Fig. S10.** Path models for testing whether increasing soil salinity indirectly enzymatic turnover via the richness difference component of Sorensen dissimilarity

**Table captions**

**Table S1.** List of the variables used in this study

**Table S2.** Summary of the global fit measures of the path model at each spatial dimension

**Table S3.** Outputs of the variance partitioning analysis

**Table S4.** Outputs of the path models

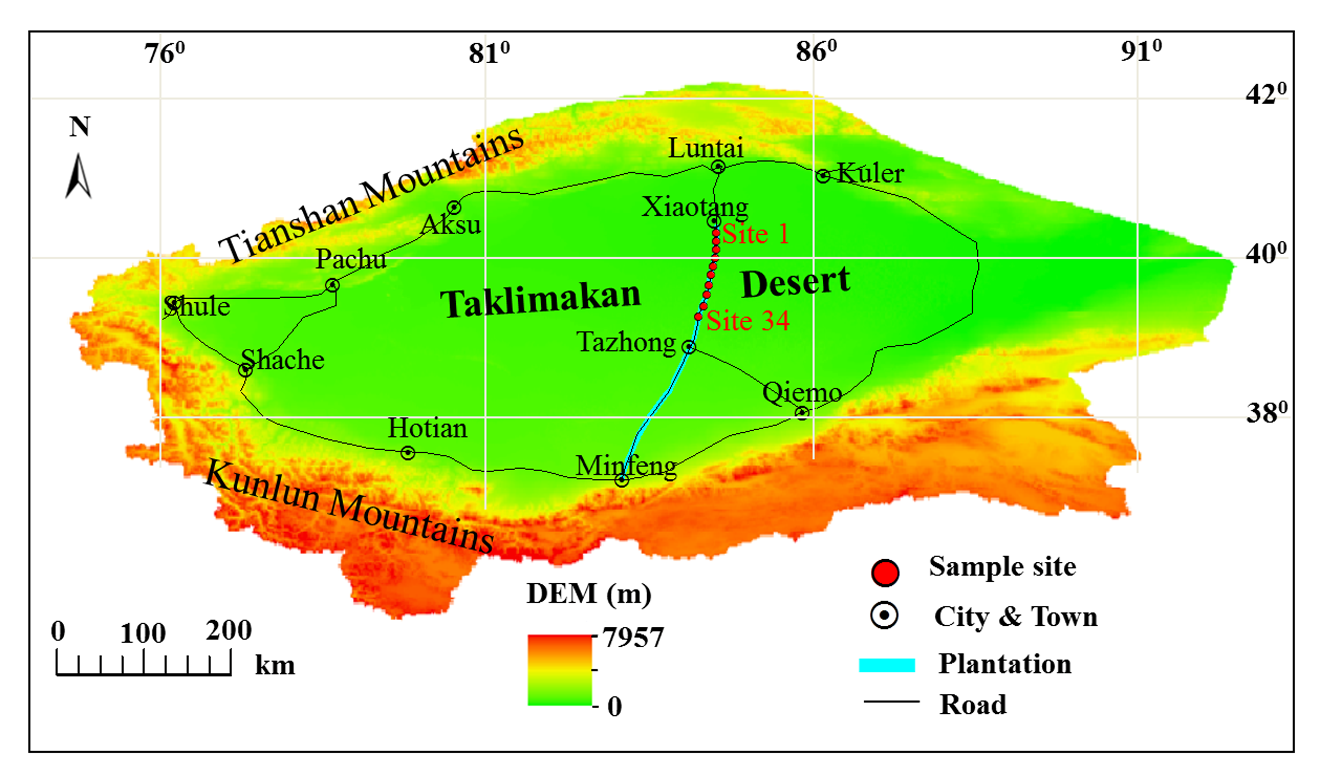
Fig. S1. Geographic location of the study sites. Sampling sites are located along the Tarim highway from Xiaotang to Tazhong.

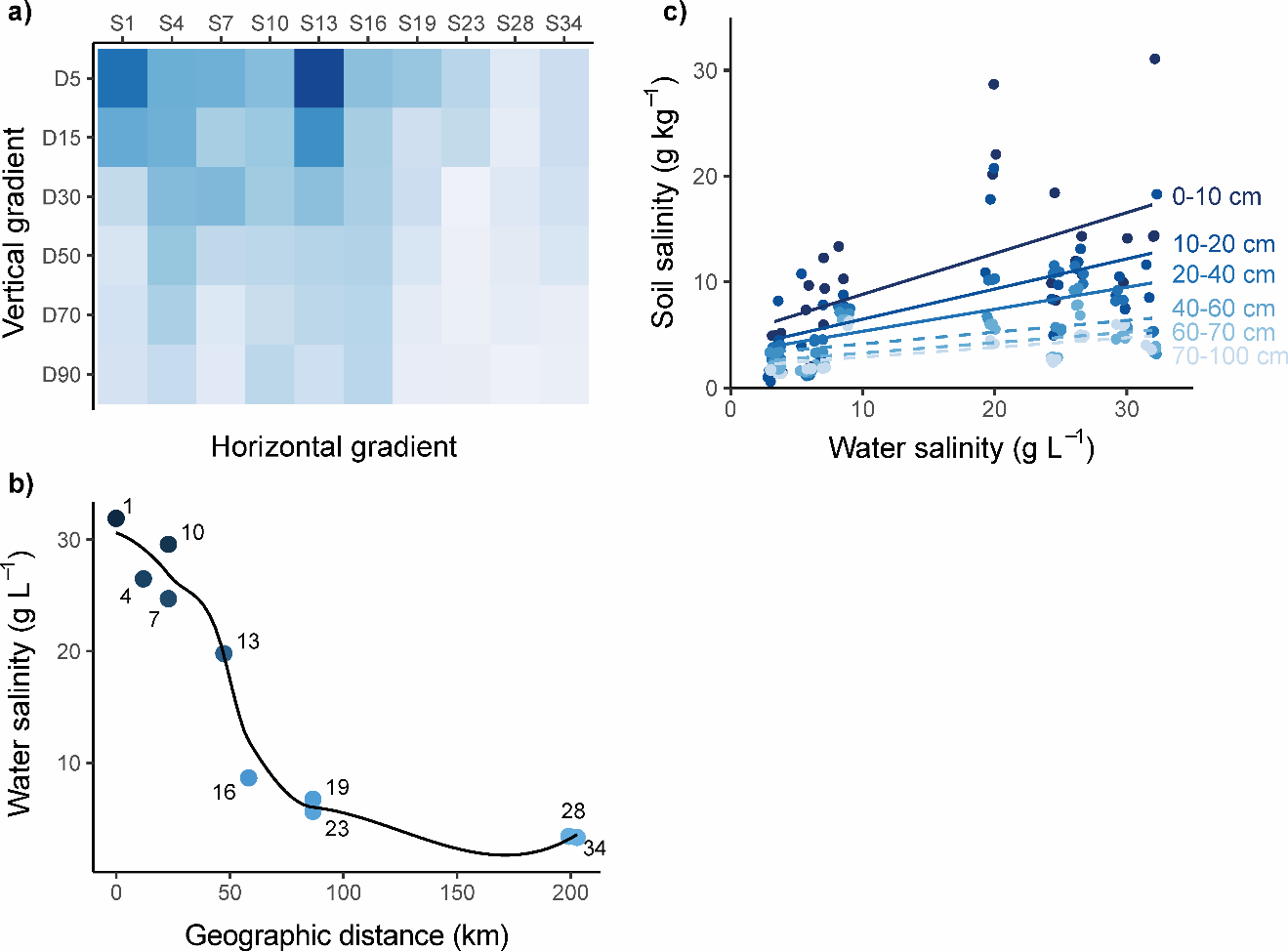

**Fig. S2.** Horizontal and vertical salinity gradients generated by saline water irrigation. a) Heatmap illustrating changes in soil salinity along the horizontal and vertical dimensions of soil depth profiles. The codes for the sampling sites are shown on the x-axis, and the average sampling depth for each soil layer is shown on the y-axis. b) The variability in irrigation water salinity along the horizontal gradient (geographic distance). Sampling sites are given next to the points. Site 1 with the highest water salinity is selected as the reference site for calculating the geographic distance (km). c) Impacts of irrigation water salinity on soil salinity across soil layers of the vertical dimension. Linear mixed-effects models are used for the significance tests. Solid lines denote significant effects of irrigation water salinity on soil salinity and dashed lines denote insignificant effects. The levels of soil/water salinity in a-c) are represented by colors from dark-blue (high soil/water salinity) to light-blue (low soil/water salinity).

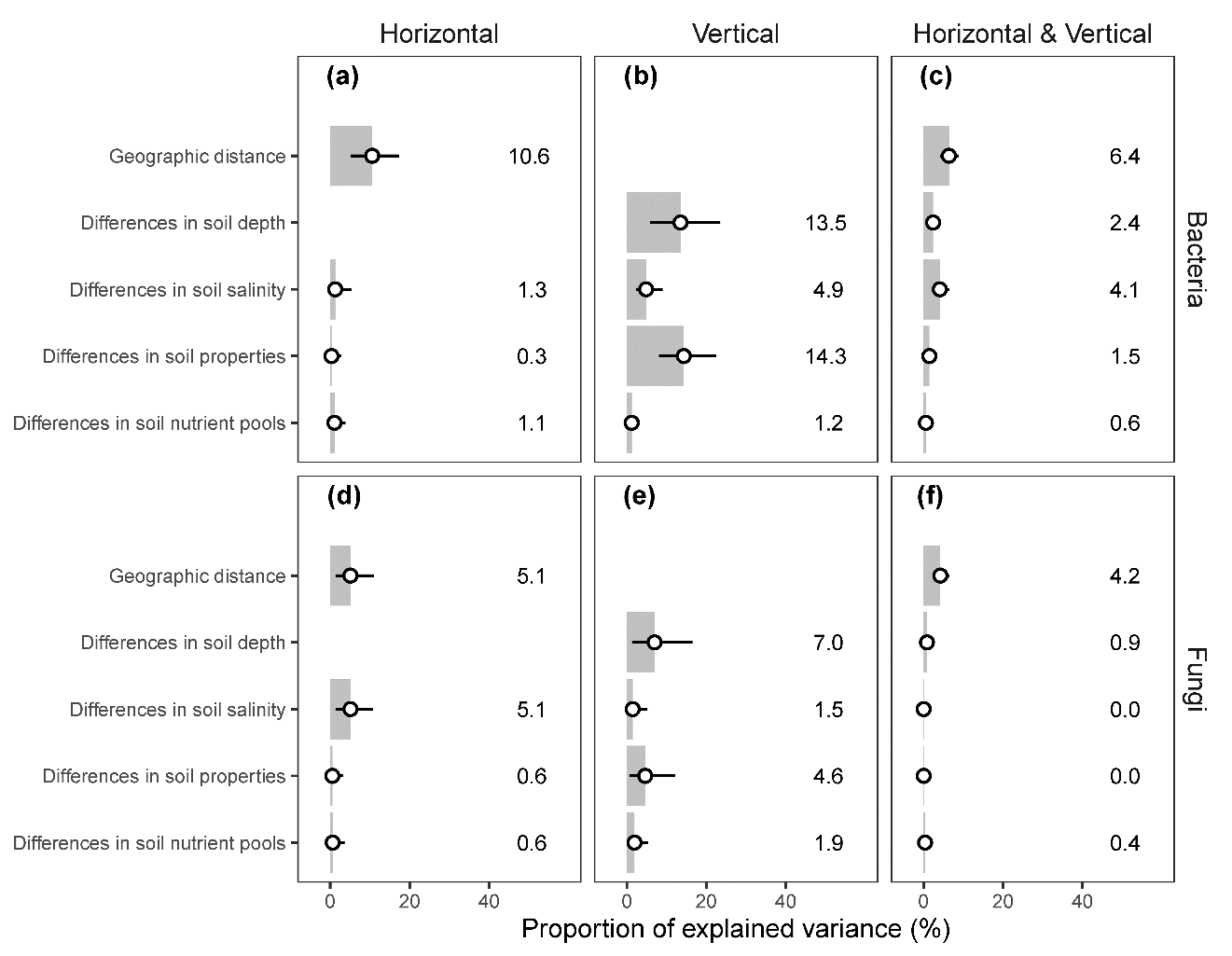

**Fig. S3.** Independent effects of explanatory variables on the replacement component of microbial β-diversity across the horizontal and vertical spatial dimensions. Bars represent the pure variance explained by the explanatory variables, and the values are shown along the bars. Points with a 95% confidence interval denote the variability of the explained variance for each explanatory variable. The replacement component of Sorensen dissimilarity index is shown. See Table S3 for values of explained variance.

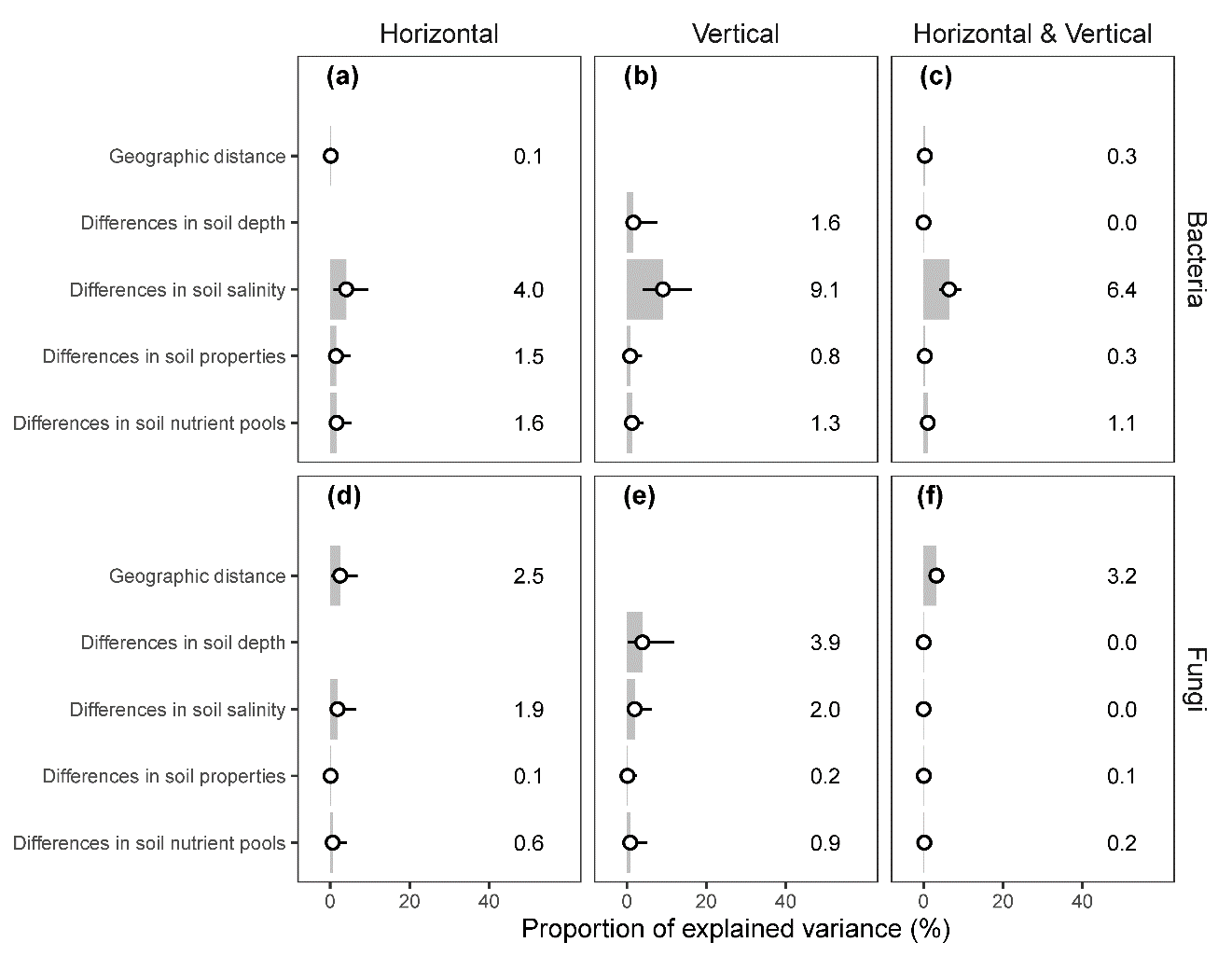

**Fig. S4.** Independent effects of explanatory variables on the richness difference component of microbial β-diversity across the horizontal and vertical spatial dimensions. Bars represent the pure variance explained by the explanatory variables, and the values are shown along the bars. Points with a 95% confidence interval denote the variability of the explained variance for each explanatory variable. The richness difference component of Sorensen dissimilarity index is shown. See Table S3 for values of explained variance.

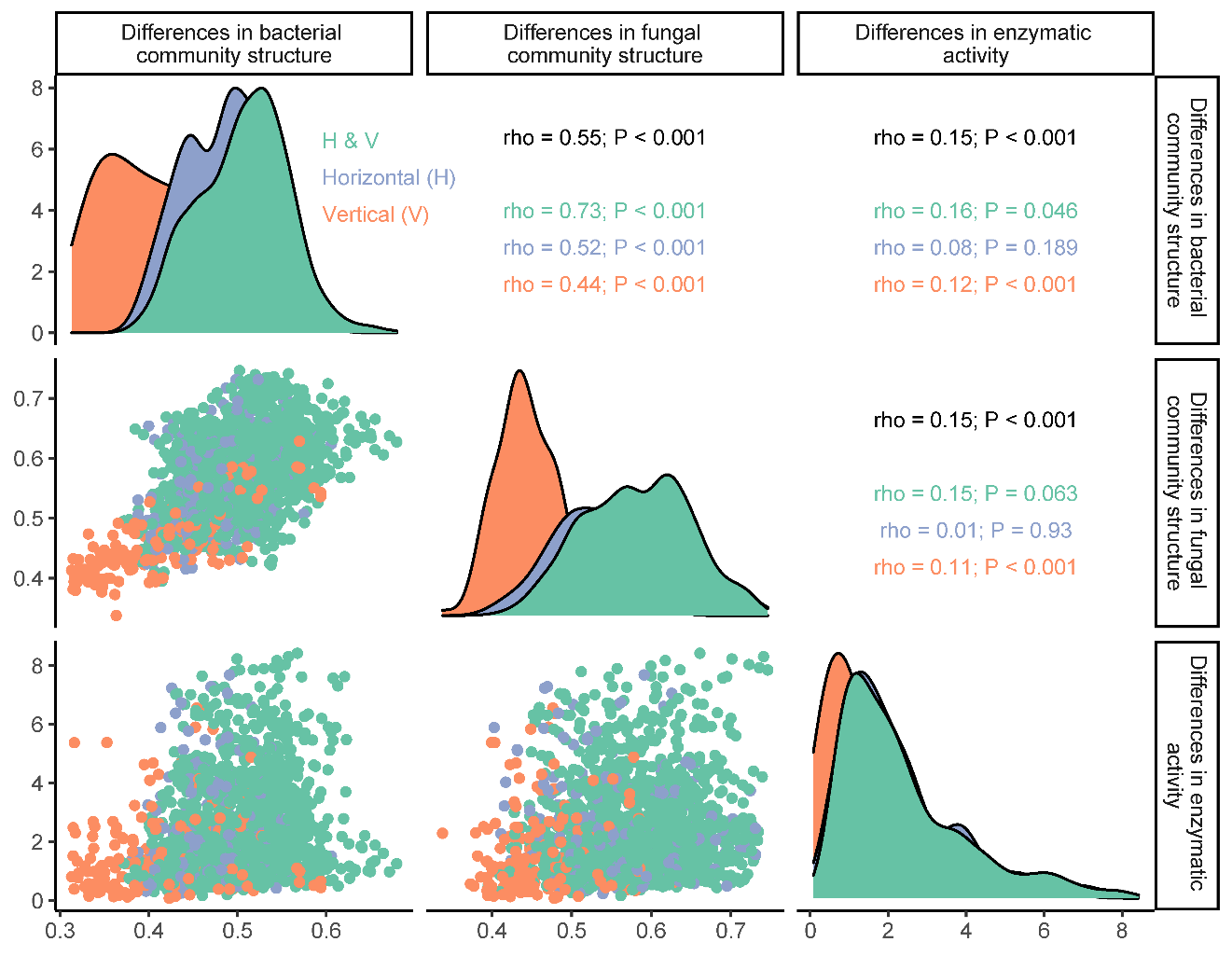

Fig. S5. Bivariate relationships between differences in soil microbial community structure and differences in enzymatic activity across spatial dimensions. Three spatial dimensions are considered including the vertical (V), horizontal (H), and horizontal & vertical (H & V) dimensions.

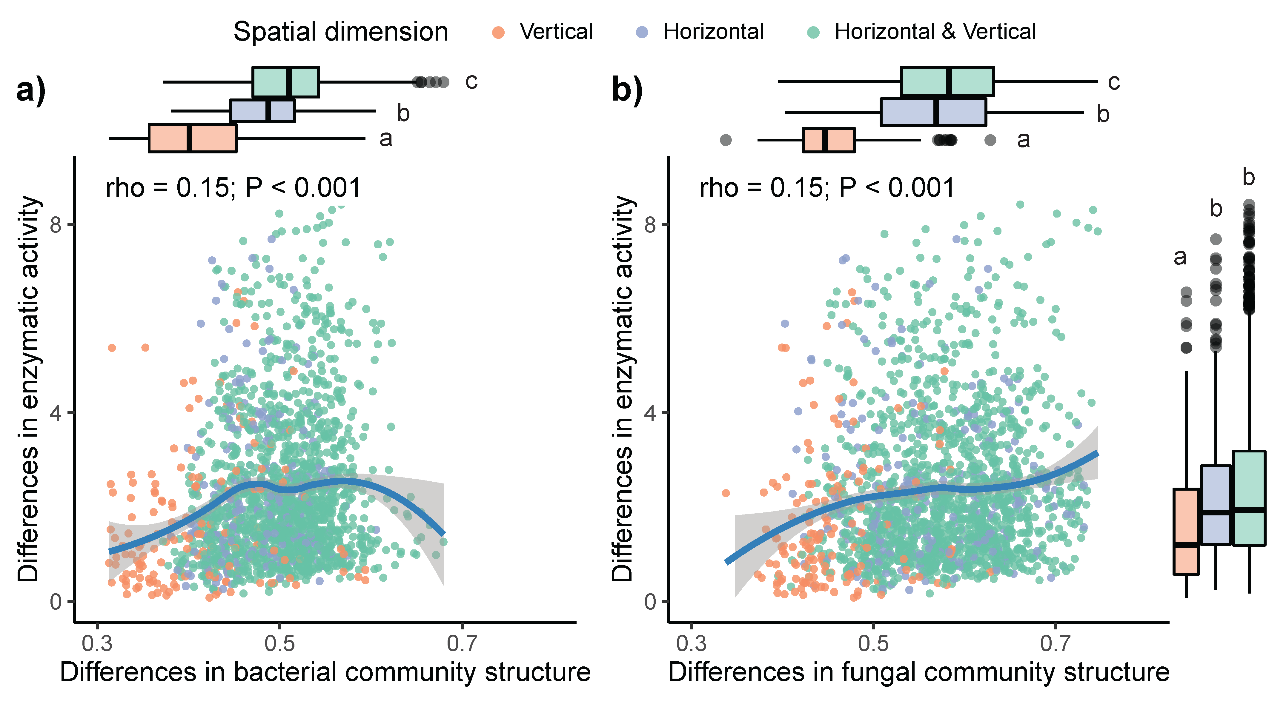

**Fig. S6.** Bivariate relationships between differences in soil microbial community structure and differences in enzymatic activity. a) bacterial community structure-enzymatic activity relationships; b) fungal community structure-enzymatic activity relationships. Lines denote the loess fitted curves showing the direction of the bivariate relationships. Shaded areas denote the 95% confidence interval. Spearman correlation coefficients are used to show the strength of the bivariate relationships. Boxplots with median, quartiles, and extreme values denote the data distributions of differences in community structure (top) and differences in enzymatic activity (right). Different letters denote significant differences among spatial dimensions.

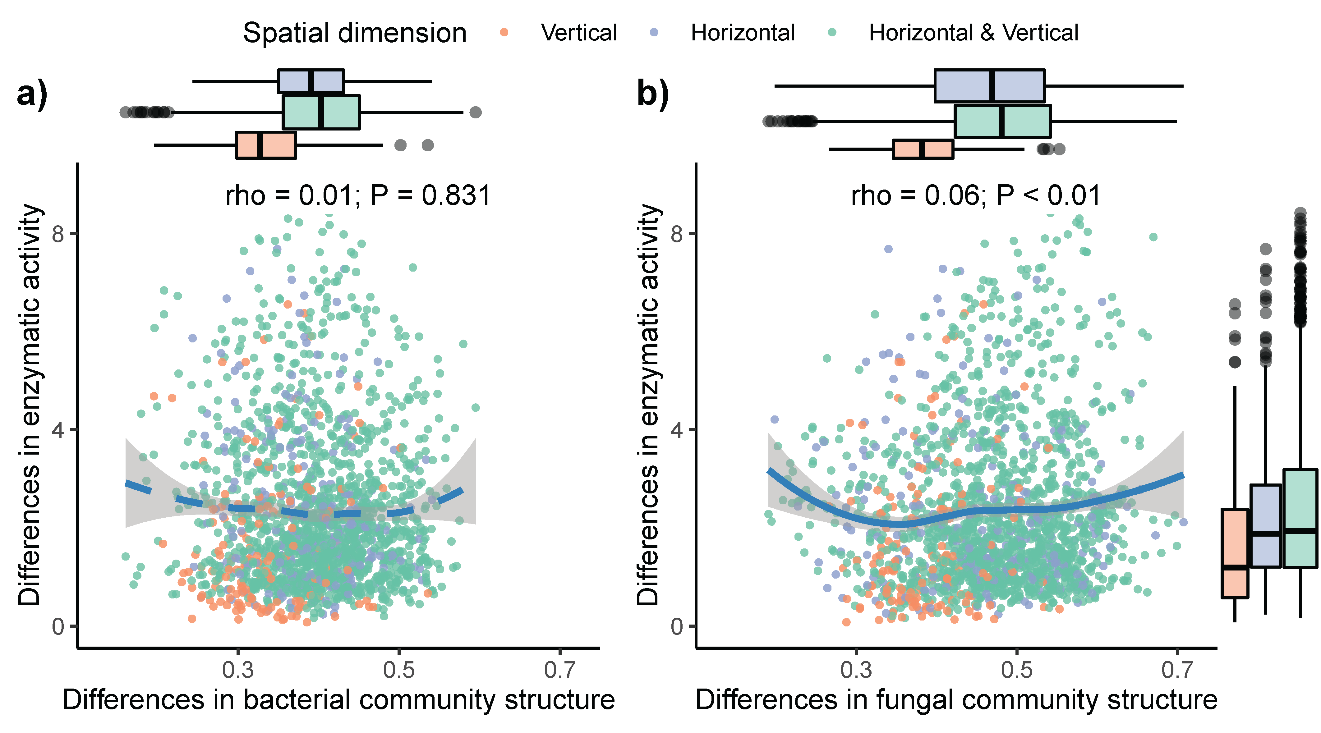

Fig. S7. Bivariate relationships between differences in the replacement component of soil microbial community structure and differences in enzymatic activity.

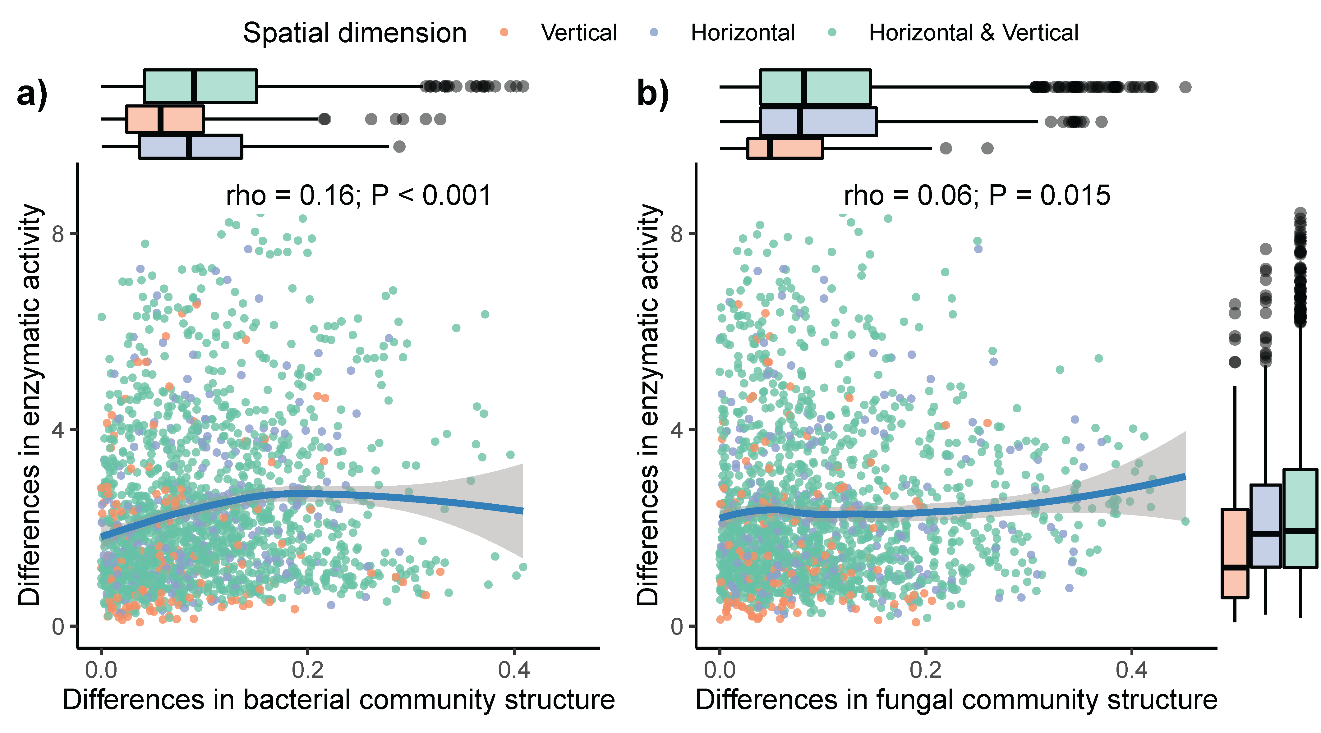

Fig. S8. Bivariate relationships between differences in the richness difference component of soil microbial community structure and differences in enzymatic activity.

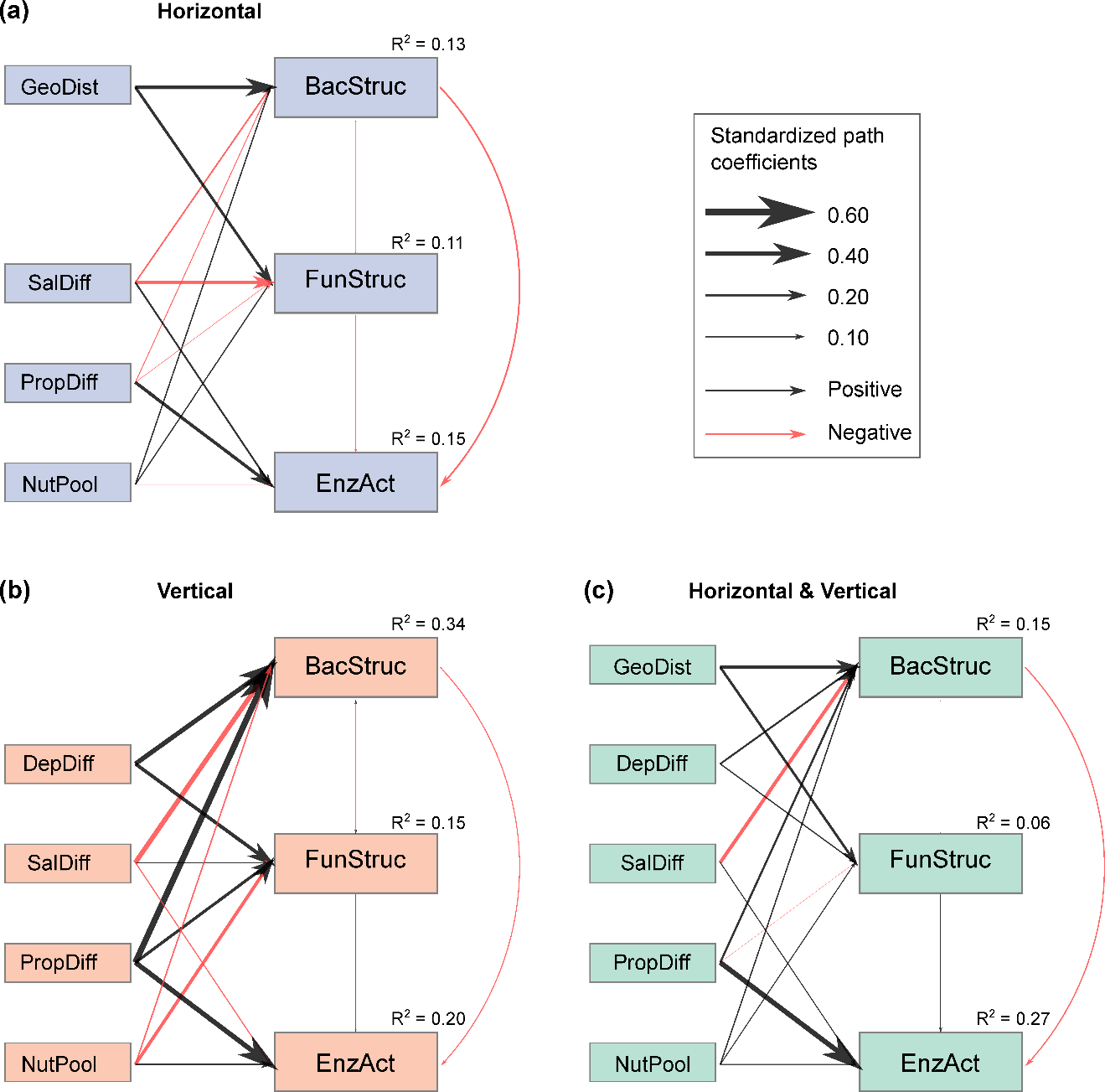

**Fig. S9.** Path models for testing whether increasing soil salinity indirectly enzymatic turnover via the replacement component of Sorensen dissimilarity. Three path models are shown based on spatial dimensions, including a) pure horizontal dimension; b) pure vertical dimension; c) combined horizontal & vertical dimension. The width of the arrows is proportional to the standardized path coefficients. Black arrows denote positive path coefficients and red arrows denote negative path coefficients. Bootstraps are used for the significance test of path coefficients (see Tables S3-S4 for details). R^2^ denotes the variance in community structure and enzymatic activity explained by environmental factors, including geographic distance (GeoDist), differences in soil depth (DepDiff), differences in soil salinity (SalDiff), differences in soil properties (PropDiff), and differences in nutrient pools (NutPool). BacStruc, FunStruc, and EnzAct denote differences in bacterial community structure, fungal community structure, and enzymatic activity, respectively.

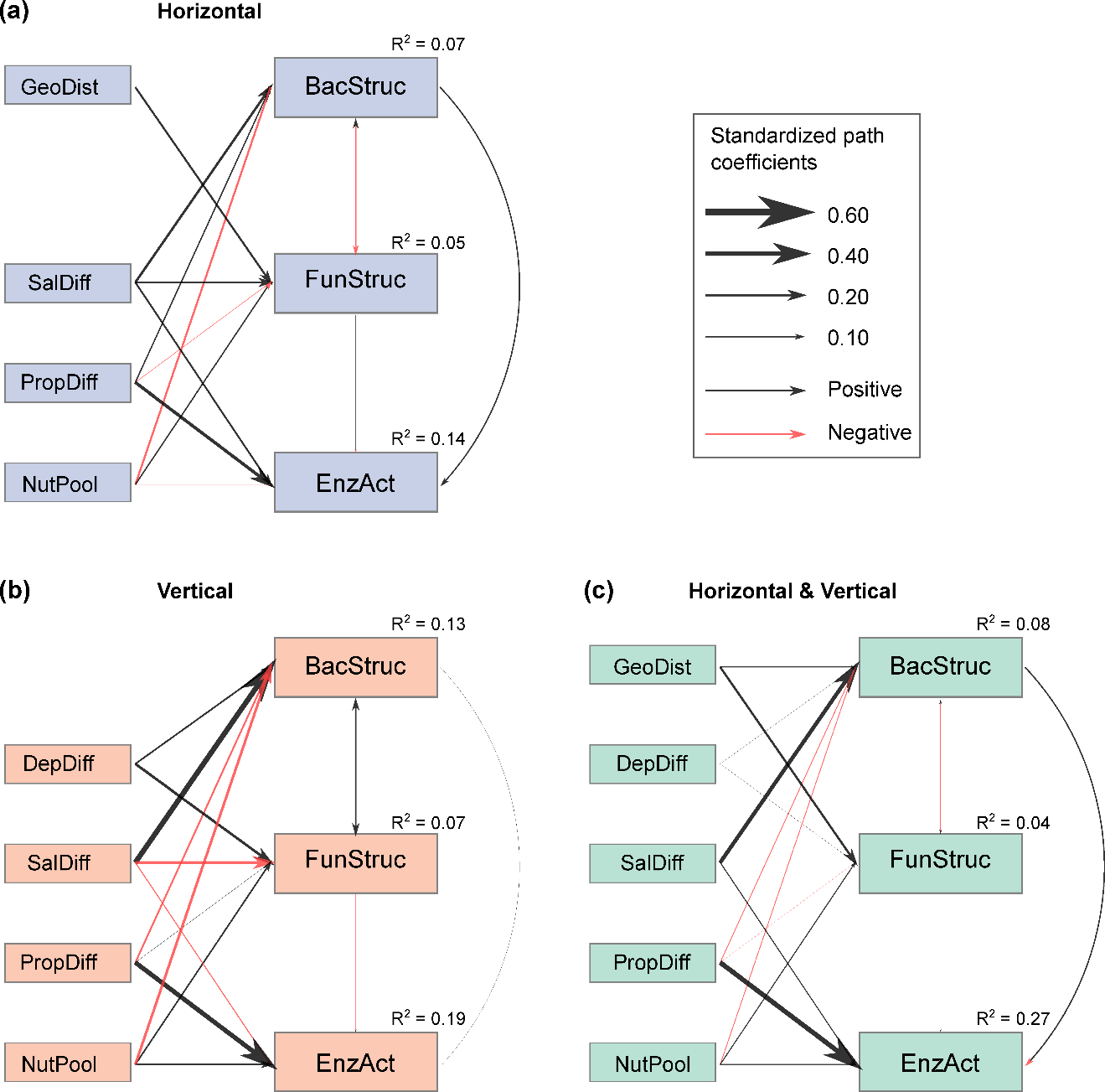

**Fig. S10.** Path models for testing whether increasing soil salinity indirectly enzymatic turnover via the richness difference component of Sorensen dissimilarity. Three path models are shown based on spatial dimensions, including a) pure horizontal dimension; b) pure vertical dimension; c) combined horizontal & vertical dimension. The width of the arrows is proportional to the standardized path coefficients. Black arrows denote positive path coefficients and red arrows denote negative path coefficients. Bootstraps are used for the significance test of path coefficients (see Tables S3-S4 for details). R^2^ denotes the variance in community structure and enzymatic activity explained by environmental factors, including geographic distance (GeoDist), differences in soil depth (DepDiff), differences in soil salinity (SalDiff), differences in soil properties (PropDiff), and differences in nutrient pools (NutPool). BacStruc, FunStruc, and EnzAct denote differences in bacterial community structure, fungal community structure, and enzymatic activity, respectively.

Table S1. List of the variables used in this study.

| Variable | Description | Unit |
| --- | --- | --- |
| Latitude | Latitude of the sampling sites used to calculate geographic distance | Degree |
| Longitude | Longitude of the sampling sites used to calculate geographic distance | Degree |
| Salinity | Salt concentration in water or in soils; soil salinity is also used to quantify salinity differences | Water salinity, g l^-1^; soil salinity, g kg^-1^ |
| pH | Soil pH that is used to quantify differences in soil properties | unitless |
| Clay | Soil clay content that is used to quantify differences in soil properties | % |
| Silt | Soil silt content that is used to quantify differences in soil properties | % |
| Sand | Soil sand content that is used to quantify differences in soil properties | % |
| Moisture | Soil gravimetric water content that is used to quantify differences in soil properties | % |
| β-glucosidase | Activity of beta-glucosidase involving in soil C cycling that is used to quantify soil microbial functional differences | nmol g^-1^ h^-1^ |
| cellobiohydrolase | Activity of cellobiohydrolase involving in soil carbon cycling that is used to quantify soil microbial functional differences | nmol g^-1^ h^-1^ |
| N-acetyl-Glucosaminidase | Activity of N-acetyl-glucosaminidase involving in soil nitrogen cycling that is used to quantify soil microbial functional differences | nmol g^-1^ h^-1^ |
| Alkaline phosphatase | Activity of alkaline phosphatase involving in soil phosphorus cycling that is used to quantify soil microbial functional differences | nmol g^-1^ h^-1^ |
| SOC | Soil organic carbon that is used to quantify soil nutrient pools | g kg^-1^ |
| STN | Soil total nitrogen that is used to quantify soil nutrient pools | g kg^-1^ |
| avaP | Soil available phosphorus that is used to quantify soil nutrient pools | g kg^-1^ |

Table S2. Summary of the global fit measures of the path model at each spatial dimension. Chisq is χ^2^, df is degree of freedom, pvalue is the P value of χ^2^ statistic test, cfi is comparative fit index, rmsea is root mean square error of approximation and srmr is standardized root mean square residual. An acceptable model fit is specified if the P value of χ^2^ statistic test > 0.05, cfi > 0.90, rmsea < 0.05 and srmr < 0.10. Note the P value of χ^2^ statistic test tends to less than 0.05 if the sample size is too large, e.g., the horizontal & vertical spatial dimension (H&V). As an alternative, the cfi, rmsea and srmr can be used to assess the model fit.

| Dimension | Component | chisq | df | pvalue | cfi | rmsea | srmr |
| --- | --- | --- | --- | --- | --- | --- | --- |
| Horizontal | β_sor_ | 0.03 | 1 | 0.860 | 1.000 | 0.000 | 0.002 |
| Vertical | β_sor_ | 2.33 | 1 | 0.127 | 0.995 | 0.094 | 0.015 |
| H&V | β_sor_ | 17.12 | 2 | <0.001 | 0.990 | 0.075 | 0.013 |
| Horizontal | β_repl_ | 0.41 | 1 | 0.520 | 1.000 | 0.000 | 0.006 |
| Vertical | β_repl_ | 2.48 | 1 | 0.115 | 0.986 | 0.099 | 0.017 |
| H&V | β_repl_ | 4.24 | 2 | 0.120 | 0.997 | 0.029 | 0.008 |
| Horizontal | β_rich_ | 0.27 | 1 | 0.606 | 1.000 | 0.000 | 0.005 |
| Vertical | β_rich_ | 1.46 | 1 | 0.227 | 0.991 | 0.055 | 0.014 |
| H&V | β_rich_ | 6.35 | 2 | 0.042 | 0.992 | 0.040 | 0.009 |

Table S3. Outputs of the variance partitioning analysis. Independent effects with the lower and upper 95% confidence interval are shown. env is differences in soil properties, geo is geographic distance, dep is differences in soil depth, nut is differences in nutrient pools, sal is differences in soil salinity. Predictor denotes the explanatory variable; Variance denotes the pure proportion of explained variance (%); lwr CI denotes the lower 95% confidence interval; upr CI denotes the upper 95% confidence interval.

| Dimension | Group | Component | Predictor | Variance | lwr CI | upr CI |
| --- | --- | --- | --- | --- | --- | --- |
| Horizontal | Bacteria | β_sor_ | env | 1.14 | 0.12 | 4.17 |
| Horizontal | Bacteria | β_sor_ | geo | 25.21 | 16.42 | 34.26 |
| Horizontal | Bacteria | β_sor_ | nut | 0.38 | 0.09 | 2.53 |
| Horizontal | Bacteria | β_sor_ | sal | 2.12 | 0.22 | 6.21 |
| Vertical | Bacteria | β_sor_ | dep | 21.95 | 12.42 | 32.78 |
| Vertical | Bacteria | β_sor_ | env | 11.05 | 4.49 | 21.28 |
| Vertical | Bacteria | β_sor_ | nut | 2.93 | 2.07 | 5.39 |
| Vertical | Bacteria | β_sor_ | sal | 3.29 | 1.43 | 6.97 |
| H&V | Bacteria | β_sor_ | dep | 5.02 | 3.24 | 7.12 |
| H&V | Bacteria | β_sor_ | env | 2.85 | 1.63 | 4.46 |
| H&V | Bacteria | β_sor_ | geo | 20.69 | 17.16 | 24.22 |
| H&V | Bacteria | β_sor_ | nut | 0.47 | 0.25 | 1.07 |
| H&V | Bacteria | β_sor_ | sal | 1.32 | 0.57 | 2.43 |
| Horizontal | Fungi | β_sor_ | env | 1.03 | 0.11 | 4.15 |
| Horizontal | Fungi | β_sor_ | geo | 24.10 | 15.26 | 33.38 |
| Horizontal | Fungi | β_sor_ | nut | 0.07 | 0.04 | 1.47 |
| Horizontal | Fungi | β_sor_ | sal | 1.70 | 0.16 | 5.55 |
| Vertical | Fungi | β_sor_ | dep | 23.75 | 12.70 | 35.82 |
| Vertical | Fungi | β_sor_ | env | 5.91 | 1.23 | 14.19 |
| Vertical | Fungi | β_sor_ | nut | 1.42 | 0.88 | 3.53 |
| Vertical | Fungi | β_sor_ | sal | 1.19 | 0.53 | 3.13 |
| H&V | Fungi | β_sor_ | dep | 0.94 | 0.26 | 2.06 |
| H&V | Fungi | β_sor_ | env | 0.27 | 0.03 | 0.99 |
| H&V | Fungi | β_sor_ | geo | 25.33 | 20.70 | 30.04 |
| H&V | Fungi | β_sor_ | nut | 0.07 | 0.01 | 0.52 |
| H&V | Fungi | β_sor_ | sal | 0.07 | 0.01 | 0.54 |
| Horizontal | Bacteria | β_repl_ | env | 0.34 | 0.05 | 2.77 |
| Horizontal | Bacteria | β_repl_ | geo | 10.58 | 5.14 | 17.23 |
| Horizontal | Bacteria | β_repl_ | nut | 1.07 | 0.06 | 3.80 |
| Horizontal | Bacteria | β_repl_ | sal | 1.26 | 0.06 | 5.39 |
| Vertical | Bacteria | β_repl_ | dep | 13.50 | 5.91 | 23.52 |
| Vertical | Bacteria | β_repl_ | env | 14.34 | 8.13 | 22.49 |
| Vertical | Bacteria | β_repl_ | nut | 1.22 | 0.82 | 2.71 |
| Vertical | Bacteria | β_repl_ | sal | 4.85 | 2.34 | 8.98 |
| H&V | Bacteria | β_repl_ | dep | 2.41 | 1.15 | 4.23 |
| H&V | Bacteria | β_repl_ | env | 1.47 | 0.63 | 2.78 |
| H&V | Bacteria | β_repl_ | geo | 6.40 | 4.26 | 8.87 |
| H&V | Bacteria | β_repl_ | nut | 0.60 | 0.20 | 1.40 |
| H&V | Bacteria | β_repl_ | sal | 4.10 | 2.25 | 6.49 |
| Horizontal | Fungi | β_repl_ | env | 0.56 | 0.09 | 3.26 |
| Horizontal | Fungi | β_repl_ | geo | 5.10 | 1.37 | 10.97 |
| Horizontal | Fungi | β_repl_ | nut | 0.64 | 0.04 | 3.58 |
| Horizontal | Fungi | β_repl_ | sal | 5.12 | 1.36 | 10.76 |
| Vertical | Fungi | β_repl_ | dep | 6.99 | 1.29 | 16.51 |
| Vertical | Fungi | β_repl_ | env | 4.63 | 0.63 | 12.22 |
| Vertical | Fungi | β_repl_ | nut | 1.93 | 0.79 | 5.25 |
| Vertical | Fungi | β_repl_ | sal | 1.47 | 0.36 | 5.16 |
| H&V | Fungi | β_repl_ | dep | 0.87 | 0.16 | 2.11 |
| H&V | Fungi | β_repl_ | env | 0.04 | 0.02 | 0.37 |
| H&V | Fungi | β_repl_ | geo | 4.25 | 2.46 | 6.45 |
| H&V | Fungi | β_repl_ | nut | 0.39 | 0.04 | 1.19 |
| H&V | Fungi | β_repl_ | sal | 0.03 | 0.01 | 0.39 |
| Horizontal | Bacteria | β_rich_ | env | 1.47 | 0.15 | 5.11 |
| Horizontal | Bacteria | β_rich_ | geo | 0.15 | 0.01 | 2.16 |
| Horizontal | Bacteria | β_rich_ | nut | 1.63 | 0.12 | 5.26 |
| Horizontal | Bacteria | β_rich_ | sal | 4.03 | 0.81 | 9.61 |
| Vertical | Bacteria | β_rich_ | dep | 1.64 | 0.13 | 7.57 |
| Vertical | Bacteria | β_rich_ | env | 0.84 | 0.39 | 3.82 |
| Vertical | Bacteria | β_rich_ | nut | 1.30 | 0.57 | 4.11 |
| Vertical | Bacteria | β_rich_ | sal | 9.09 | 4.00 | 16.28 |
| H&V | Bacteria | β_rich_ | dep | 0.02 | 0.01 | 0.39 |
| H&V | Bacteria | β_rich_ | env | 0.33 | 0.21 | 0.72 |
| H&V | Bacteria | β_rich_ | geo | 0.34 | 0.01 | 1.19 |
| H&V | Bacteria | β_rich_ | nut | 1.09 | 0.44 | 2.17 |
| H&V | Bacteria | β_rich_ | sal | 6.45 | 3.92 | 9.46 |
| Horizontal | Fungi | β_rich_ | env | 0.10 | 0.04 | 2.04 |
| Horizontal | Fungi | β_rich_ | geo | 2.50 | 0.30 | 6.88 |
| Horizontal | Fungi | β_rich_ | nut | 0.64 | 0.03 | 4.20 |
| Horizontal | Fungi | β_rich_ | sal | 1.86 | 0.06 | 6.49 |
| Vertical | Fungi | β_rich_ | dep | 3.91 | 0.22 | 11.95 |
| Vertical | Fungi | β_rich_ | env | 0.16 | 0.10 | 2.52 |
| Vertical | Fungi | β_rich_ | nut | 0.87 | 0.14 | 5.18 |
| Vertical | Fungi | β_rich_ | sal | 1.96 | 0.27 | 6.24 |
| H&V | Fungi | β_rich_ | dep | 0.04 | 0.00 | 0.52 |
| H&V | Fungi | β_rich_ | env | 0.08 | 0.01 | 0.54 |
| H&V | Fungi | β_rich_ | geo | 3.23 | 1.75 | 5.19 |
| H&V | Fungi | β_rich_ | nut | 0.20 | 0.01 | 0.81 |
| H&V | Fungi | β_rich_ | sal | 0.03 | 0.01 | 0.43 |

Table S4. Outputs of the path models. Resp = response variable; op = operator type (~ denotes regression and ~~ denotes covariance); Pred = predictor variable; β = raw estimate of path coefficient; SE = standard error; Z = Z statistic test; P = P value of the estimate; β_std_ = standardized estimate of path coefficient. β_sor_ is the Sorensen dissimilarity index, β_repl_ is the replacement component of Sorensen dissimilarity index, β_rich_ is the richness difference component of Sorensen dissimilarity index, enz is differences in enzymatic activity, geo is the geographic distance, dep is differences in soil depth, sal is differences in soil salinity, env is differences in soil properties, and nut is differences in soil nutrient pools.

| Dimension | Component | Resp | op | Pred | β | SE | Z | P | β_std_ |
| --- | --- | --- | --- | --- | --- | --- | --- | --- | --- |
| Horizontal | β_sor_ | bac.sor | ~ | geo | 0.039 | 0.004 | 10.19 | 0.000 | 0.50 |
| Horizontal | β_sor_ | bac.sor | ~ | sal | 0.004 | 0.002 | 2.30 | 0.022 | 0.13 |
| Horizontal | β_sor_ | bac.sor | ~ | env | 0.003 | 0.003 | 1.04 | 0.298 | 0.06 |
| Horizontal | β_sor_ | bac.sor | ~ | nut | -0.004 | 0.002 | -1.66 | 0.096 | -0.10 |
| Horizontal | β_sor_ | fun.sor | ~ | geo | 0.064 | 0.007 | 9.36 | 0.000 | 0.50 |
| Horizontal | β_sor_ | fun.sor | ~ | sal | -0.006 | 0.003 | -2.27 | 0.023 | -0.13 |
| Horizontal | β_sor_ | fun.sor | ~ | env | -0.008 | 0.005 | -1.73 | 0.084 | -0.10 |
| Horizontal | β_sor_ | fun.sor | ~ | nut | 0.000 | 0.003 | -0.08 | 0.938 | 0.00 |
| Horizontal | β_sor_ | enz | ~ | bac.sor | -1.616 | 2.419 | -0.67 | 0.504 | -0.05 |
| Horizontal | β_sor_ | enz | ~ | fun.sor | -0.158 | 1.451 | -0.11 | 0.914 | -0.01 |
| Horizontal | β_sor_ | enz | ~ | sal | 0.163 | 0.054 | 3.04 | 0.002 | 0.17 |
| Horizontal | β_sor_ | enz | ~ | env | 0.447 | 0.123 | 3.64 | 0.000 | 0.28 |
| Horizontal | β_sor_ | enz | ~ | nut | -0.050 | 0.067 | -0.74 | 0.462 | -0.04 |
| Horizontal | β_sor_ | bac.sor | ~~ | fun.sor | 0.001 | 0.000 | 7.30 | 0.000 | 0.42 |
| Vertical | β_sor_ | bac.sor | ~ | dep | 0.002 | 0.000 | 6.46 | 0.000 | 0.51 |
| Vertical | β_sor_ | bac.sor | ~ | sal | 0.003 | 0.003 | 0.92 | 0.358 | 0.07 |
| Vertical | β_sor_ | bac.sor | ~ | env | 0.015 | 0.003 | 4.42 | 0.000 | 0.37 |
| Vertical | β_sor_ | bac.sor | ~ | nut | -0.018 | 0.004 | -4.63 | 0.000 | -0.32 |
| Vertical | β_sor_ | fun.sor | ~ | dep | 0.001 | 0.000 | 6.38 | 0.000 | 0.55 |
| Vertical | β_sor_ | fun.sor | ~ | sal | -0.005 | 0.002 | -2.08 | 0.038 | -0.17 |
| Vertical | β_sor_ | fun.sor | ~ | env | 0.010 | 0.003 | 3.16 | 0.002 | 0.30 |
| Vertical | β_sor_ | fun.sor | ~ | nut | -0.008 | 0.003 | -2.31 | 0.021 | -0.17 |
| Vertical | β_sor_ | enz | ~ | bac.sor | -2.170 | 2.517 | -0.86 | 0.389 | -0.10 |
| Vertical | β_sor_ | enz | ~ | fun.sor | 1.874 | 3.314 | 0.57 | 0.572 | 0.07 |
| Vertical | β_sor_ | enz | ~ | sal | -0.022 | 0.106 | -0.21 | 0.832 | -0.03 |
| Vertical | β_sor_ | enz | ~ | env | 0.368 | 0.112 | 3.29 | 0.001 | 0.41 |
| Vertical | β_sor_ | enz | ~ | nut | 0.136 | 0.139 | 0.98 | 0.326 | 0.11 |
| Vertical | β_sor_ | bac.sor | ~~ | fun.sor | 0.001 | 0.000 | 5.08 | 0.000 | 0.67 |
| H&V | β_sor_ | bac.sor | ~ | geo | 0.040 | 0.002 | 21.29 | 0.000 | 0.46 |
| H&V | β_sor_ | bac.sor | ~ | dep | 0.000 | 0.000 | 8.93 | 0.000 | 0.22 |
| H&V | β_sor_ | bac.sor | ~ | sal | 0.002 | 0.001 | 3.33 | 0.001 | 0.09 |
| H&V | β_sor_ | bac.sor | ~ | env | 0.007 | 0.001 | 6.35 | 0.000 | 0.17 |
| H&V | β_sor_ | bac.sor | ~ | nut | -0.005 | 0.001 | -4.23 | 0.000 | -0.12 |
| H&V | β_sor_ | fun.sor | ~ | geo | 0.059 | 0.003 | 19.31 | 0.000 | 0.50 |
| H&V | β_sor_ | fun.sor | ~ | dep | 0.000 | 0.000 | 4.08 | 0.000 | 0.10 |
| H&V | β_sor_ | fun.sor | ~ | sal | 0.001 | 0.001 | 1.22 | 0.223 | 0.04 |
| H&V | β_sor_ | fun.sor | ~ | env | -0.004 | 0.001 | -2.45 | 0.014 | -0.07 |
| H&V | β_sor_ | fun.sor | ~ | nut | 0.002 | 0.002 | 1.01 | 0.310 | 0.03 |
| H&V | β_sor_ | enz | ~ | bac.sor | -1.357 | 0.883 | -1.54 | 0.124 | -0.04 |
| H&V | β_sor_ | enz | ~ | fun.sor | 3.143 | 0.654 | 4.81 | 0.000 | 0.13 |
| H&V | β_sor_ | enz | ~ | sal | 0.084 | 0.024 | 3.46 | 0.001 | 0.10 |
| H&V | β_sor_ | enz | ~ | env | 0.542 | 0.044 | 12.26 | 0.000 | 0.44 |
| H&V | β_sor_ | enz | ~ | nut | 0.065 | 0.034 | 1.89 | 0.059 | 0.05 |
| H&V | β_sor_ | bac.sor | ~~ | fun.sor | 0.001 | 0.000 | 10.96 | 0.000 | 0.28 |
| Horizontal | β_repl_ | bac.repl | ~ | geo | 0.035 | 0.005 | 6.42 | 0.000 | 0.33 |
| Horizontal | β_repl_ | bac.repl | ~ | sal | -0.005 | 0.003 | -1.74 | 0.082 | -0.12 |
| Horizontal | β_repl_ | bac.repl | ~ | env | -0.004 | 0.004 | -0.89 | 0.373 | -0.06 |
| Horizontal | β_repl_ | bac.repl | ~ | nut | 0.006 | 0.003 | 2.00 | 0.046 | 0.10 |
| Horizontal | β_repl_ | fun.repl | ~ | geo | 0.039 | 0.009 | 4.19 | 0.000 | 0.23 |
| Horizontal | β_repl_ | fun.repl | ~ | sal | -0.015 | 0.003 | -4.19 | 0.000 | -0.24 |
| Horizontal | β_repl_ | fun.repl | ~ | env | -0.004 | 0.007 | -0.59 | 0.554 | -0.04 |
| Horizontal | β_repl_ | fun.repl | ~ | nut | 0.008 | 0.006 | 1.42 | 0.155 | 0.09 |
| Horizontal | β_repl_ | enz | ~ | bac.repl | -3.195 | 1.432 | -2.23 | 0.026 | -0.13 |
| Horizontal | β_repl_ | enz | ~ | fun.repl | -0.718 | 0.899 | -0.80 | 0.424 | -0.05 |
| Horizontal | β_repl_ | enz | ~ | sal | 0.133 | 0.054 | 2.46 | 0.014 | 0.14 |
| Horizontal | β_repl_ | enz | ~ | env | 0.432 | 0.118 | 3.66 | 0.000 | 0.27 |
| Horizontal | β_repl_ | enz | ~ | nut | -0.016 | 0.069 | -0.23 | 0.821 | -0.01 |
| Horizontal | β_repl_ | bac.repl | ~~ | fun.repl | 0.000 | 0.000 | -0.26 | 0.795 | -0.01 |
| Vertical | β_repl_ | bac.repl | ~ | dep | 0.001 | 0.000 | 5.06 | 0.000 | 0.41 |
| Vertical | β_repl_ | bac.repl | ~ | sal | -0.014 | 0.002 | -5.77 | 0.000 | -0.42 |
| Vertical | β_repl_ | bac.repl | ~ | env | 0.021 | 0.003 | 6.83 | 0.000 | 0.54 |
| Vertical | β_repl_ | bac.repl | ~ | nut | -0.006 | 0.004 | -1.48 | 0.138 | -0.11 |
| Vertical | β_repl_ | fun.repl | ~ | dep | 0.001 | 0.000 | 3.15 | 0.002 | 0.30 |
| Vertical | β_repl_ | fun.repl | ~ | sal | 0.002 | 0.003 | 0.90 | 0.369 | 0.08 |
| Vertical | β_repl_ | fun.repl | ~ | env | 0.009 | 0.003 | 2.60 | 0.009 | 0.25 |
| Vertical | β_repl_ | fun.repl | ~ | nut | -0.014 | 0.004 | -3.20 | 0.001 | -0.28 |
| Vertical | β_repl_ | enz | ~ | bac.repl | -1.712 | 2.182 | -0.78 | 0.433 | -0.07 |
| Vertical | β_repl_ | enz | ~ | fun.repl | 0.770 | 2.234 | 0.34 | 0.730 | 0.03 |
| Vertical | β_repl_ | enz | ~ | sal | -0.062 | 0.108 | -0.58 | 0.564 | -0.08 |
| Vertical | β_repl_ | enz | ~ | env | 0.383 | 0.117 | 3.27 | 0.001 | 0.43 |
| Vertical | β_repl_ | enz | ~ | nut | 0.165 | 0.140 | 1.18 | 0.240 | 0.13 |
| Vertical | β_repl_ | bac.repl | ~~ | fun.repl | 0.000 | 0.000 | -1.18 | 0.238 | -0.11 |
| H&V | β_repl_ | bac.repl | ~ | geo | 0.032 | 0.003 | 10.59 | 0.000 | 0.26 |
| H&V | β_repl_ | bac.repl | ~ | dep | 0.000 | 0.000 | 5.51 | 0.000 | 0.15 |
| H&V | β_repl_ | bac.repl | ~ | sal | -0.011 | 0.001 | -8.78 | 0.000 | -0.30 |
| H&V | β_repl_ | bac.repl | ~ | env | 0.010 | 0.002 | 5.25 | 0.000 | 0.17 |
| H&V | β_repl_ | bac.repl | ~ | nut | 0.006 | 0.002 | 3.42 | 0.001 | 0.10 |
| H&V | β_repl_ | fun.repl | ~ | geo | 0.031 | 0.004 | 8.09 | 0.000 | 0.21 |
| H&V | β_repl_ | fun.repl | ~ | dep | 0.000 | 0.000 | 3.23 | 0.001 | 0.09 |
| H&V | β_repl_ | fun.repl | ~ | sal | 0.000 | 0.001 | -0.03 | 0.979 | 0.00 |
| H&V | β_repl_ | fun.repl | ~ | env | -0.002 | 0.002 | -1.07 | 0.286 | -0.03 |
| H&V | β_repl_ | fun.repl | ~ | nut | 0.005 | 0.002 | 2.24 | 0.025 | 0.07 |
| H&V | β_repl_ | enz | ~ | bac.repl | -2.038 | 0.584 | -3.49 | 0.000 | -0.09 |
| H&V | β_repl_ | enz | ~ | fun.repl | 1.082 | 0.432 | 2.50 | 0.012 | 0.06 |
| H&V | β_repl_ | enz | ~ | sal | 0.062 | 0.025 | 2.47 | 0.014 | 0.07 |
| H&V | β_repl_ | enz | ~ | env | 0.543 | 0.043 | 12.63 | 0.000 | 0.44 |
| H&V | β_repl_ | enz | ~ | nut | 0.085 | 0.035 | 2.41 | 0.016 | 0.06 |
| H&V | β_repl_ | bac.repl | ~~ | fun.repl | 0.000 | 0.000 | -0.52 | 0.603 | -0.01 |
| Horizontal | β_rich_ | bac.rich | ~ | geo | 0.004 | 0.006 | 0.66 | 0.509 | 0.04 |
| Horizontal | β_rich_ | bac.rich | ~ | sal | 0.008 | 0.003 | 3.04 | 0.002 | 0.20 |
| Horizontal | β_rich_ | bac.rich | ~ | env | 0.007 | 0.005 | 1.49 | 0.137 | 0.10 |
| Horizontal | β_rich_ | bac.rich | ~ | nut | -0.010 | 0.003 | -2.79 | 0.005 | -0.16 |
| Horizontal | β_rich_ | fun.rich | ~ | geo | 0.025 | 0.009 | 2.96 | 0.003 | 0.16 |
| Horizontal | β_rich_ | fun.rich | ~ | sal | 0.009 | 0.004 | 2.20 | 0.028 | 0.16 |
| Horizontal | β_rich_ | fun.rich | ~ | env | -0.004 | 0.007 | -0.63 | 0.529 | -0.04 |
| Horizontal | β_rich_ | fun.rich | ~ | nut | -0.008 | 0.006 | -1.40 | 0.163 | -0.10 |
| Horizontal | β_rich_ | enz | ~ | bac.rich | 2.170 | 1.477 | 1.47 | 0.142 | 0.10 |
| Horizontal | β_rich_ | enz | ~ | fun.rich | 0.595 | 0.959 | 0.62 | 0.535 | 0.04 |
| Horizontal | β_rich_ | enz | ~ | sal | 0.134 | 0.053 | 2.53 | 0.011 | 0.14 |
| Horizontal | β_rich_ | enz | ~ | env | 0.426 | 0.122 | 3.50 | 0.000 | 0.27 |
| Horizontal | β_rich_ | enz | ~ | nut | -0.023 | 0.069 | -0.33 | 0.742 | -0.02 |
| Horizontal | β_rich_ | bac.rich | ~~ | fun.rich | -0.001 | 0.000 | -1.54 | 0.123 | -0.10 |
| Vertical | β_rich_ | bac.rich | ~ | dep | 0.000 | 0.000 | 1.46 | 0.143 | 0.13 |
| Vertical | β_rich_ | bac.rich | ~ | sal | 0.016 | 0.003 | 5.16 | 0.000 | 0.46 |
| Vertical | β_rich_ | bac.rich | ~ | env | -0.006 | 0.005 | -1.22 | 0.224 | -0.14 |
| Vertical | β_rich_ | bac.rich | ~ | nut | -0.012 | 0.005 | -2.40 | 0.016 | -0.22 |
| Vertical | β_rich_ | fun.rich | ~ | dep | 0.001 | 0.000 | 2.23 | 0.025 | 0.22 |
| Vertical | β_rich_ | fun.rich | ~ | sal | -0.007 | 0.003 | -2.58 | 0.010 | -0.24 |
| Vertical | β_rich_ | fun.rich | ~ | env | 0.001 | 0.003 | 0.28 | 0.783 | 0.03 |
| Vertical | β_rich_ | fun.rich | ~ | nut | 0.006 | 0.005 | 1.08 | 0.282 | 0.12 |
| Vertical | β_rich_ | enz | ~ | bac.rich | 0.537 | 1.720 | 0.31 | 0.755 | 0.02 |
| Vertical | β_rich_ | enz | ~ | fun.rich | -0.856 | 2.181 | -0.39 | 0.695 | -0.03 |
| Vertical | β_rich_ | enz | ~ | sal | -0.053 | 0.110 | -0.48 | 0.631 | -0.07 |
| Vertical | β_rich_ | enz | ~ | env | 0.357 | 0.105 | 3.40 | 0.001 | 0.40 |
| Vertical | β_rich_ | enz | ~ | nut | 0.169 | 0.142 | 1.19 | 0.235 | 0.13 |
| Vertical | β_rich_ | bac.rich | ~~ | fun.rich | 0.000 | 0.000 | 1.47 | 0.141 | 0.13 |
| H&V | β_rich_ | bac.rich | ~ | geo | 0.008 | 0.003 | 2.21 | 0.027 | 0.06 |
| H&V | β_rich_ | bac.rich | ~ | dep | 0.000 | 0.000 | 0.08 | 0.938 | 0.00 |
| H&V | β_rich_ | bac.rich | ~ | sal | 0.014 | 0.002 | 8.97 | 0.000 | 0.34 |
| H&V | β_rich_ | bac.rich | ~ | env | -0.003 | 0.002 | -1.42 | 0.154 | -0.05 |
| H&V | β_rich_ | bac.rich | ~ | nut | -0.011 | 0.002 | -5.20 | 0.000 | -0.16 |
| H&V | β_rich_ | fun.rich | ~ | geo | 0.027 | 0.004 | 7.57 | 0.000 | 0.18 |
| H&V | β_rich_ | fun.rich | ~ | dep | 0.000 | 0.000 | -0.51 | 0.611 | -0.01 |
| H&V | β_rich_ | fun.rich | ~ | sal | 0.001 | 0.002 | 0.77 | 0.442 | 0.03 |
| H&V | β_rich_ | fun.rich | ~ | env | -0.001 | 0.002 | -0.74 | 0.461 | -0.02 |
| H&V | β_rich_ | fun.rich | ~ | nut | -0.004 | 0.002 | -1.60 | 0.109 | -0.05 |
| H&V | β_rich_ | enz | ~ | bac.rich | 2.018 | 0.535 | 3.77 | 0.000 | 0.10 |
| H&V | β_rich_ | enz | ~ | fun.rich | 0.648 | 0.404 | 1.60 | 0.109 | 0.03 |
| H&V | β_rich_ | enz | ~ | sal | 0.056 | 0.026 | 2.19 | 0.028 | 0.07 |
| H&V | β_rich_ | enz | ~ | env | 0.528 | 0.042 | 12.51 | 0.000 | 0.42 |
| H&V | β_rich_ | enz | ~ | nut | 0.101 | 0.036 | 2.84 | 0.005 | 0.07 |
| H&V | β_rich_ | bac.rich | ~~ | fun.rich | 0.000 | 0.000 | -1.82 | 0.069 | -0.05 |
